# ER stress-associated IRE1α signalling links impaired QPRT-dependent NAD⁺ biosynthesis to reduced oocyte competence

**DOI:** 10.64898/2026.11.04.764453

**Authors:** Mohd Athar, E Ajith Kumar, Pravin Birajdar, M Rajender, Akshay Kumar, Anjali Kumari, Sahina Sabnam, Ankita Verma, L Pradnya, Partha Sarathi Nial, G. Taru Sharma, H.B.D. Prasada Rao

**Affiliations:** BRIC-National Institute of Animal Biotechnology, Hyderabad, Telangana 500032, India; Regional Centre for Biotechnology, Faridabad - 121 001, India

**Keywords:** Oocyte quality, NAD+, metabolism, Endoplasmic reticulum stress, QPRT, IRE1α-RIDD

## Abstract

Oocyte quality is closely linked to metabolic and organelle homeostasis, but the mechanisms underlying naturally occurring differences in oocyte quality remain incompletely understood. Here, we identify a link between endoplasmic reticulum (ER) stress, impaired NAD⁺ biosynthesis and poor oocyte quality. Poor-grade goat oocytes exhibited profound NAD⁺ depletion, accompanied by quinolinic acid accumulation and reduced expression of quinolinate phosphoribosyltransferase (QPRT), suggesting a bottleneck in de novo NAD⁺ biosynthesis. Restoring QPRT expression or bypassing this metabolic step improved meiotic maturation. Further analyses revealed altered ER stress signalling, increased IRE1α-associated activity and reduced QPRT RNA stability. Recombinant IRE1α processed QPRT RNA in vitro, suggesting a possible link between IRE1α activity and QPRT transcript regulation. Supplementation with nicotinamide mononucleotide (NMN) restored NAD⁺ levels and improved several features of poor-quality oocytes, including calcium homeostasis, oxidative stress and mitochondrial membrane potential. NMN treatment also improved early embryonic development, with its beneficial effects involving SIRT6 activity and changes in DNA methylation-associated fluorescence. Importantly, NMN supplementation improved meiotic maturation in poor-grade bovine and porcine oocytes, indicating that this metabolic intervention may have broader relevance across mammals. Analysis of published human oocyte proteomic data further revealed age-associated changes in ER proteostasis and NAD-related proteins. Together, our findings identify ER stress-associated disruption of QPRT-dependent NAD⁺ biosynthesis as a potential contributor to poor oocyte quality and suggest that restoring NAD⁺ metabolism may offer a strategy to improve the functional properties of compromised oocytes.

## Introduction

Oocyte developmental competence is a major determinant of fertilization, embryo development and female reproductive success ^1,2^. In assisted reproduction and livestock embryo production, oocytes are routinely selected according to morphological features such as the extent of cumulus-cell association, cytoplasmic appearance and meiotic status ^3,4^. Oocytes surrounded by compact, multi layered cumulus cells and displaying homogeneous cytoplasm are generally classified as excellent- or high-grade, whereas oocytes with sparse or fragmented cumulus investment or altered cytoplasmic morphology are considered poor-grade^4–8^. These criteria are useful for selecting oocytes with a higher probability of developmental success, but they provide little information about the cellular state underlying this difference in competence. Oocyte heterogeneity is evident at the moment of follicular collection. Oocytes drawn from the same ovarian population can differ markedly in morphology and developmental potential, even though none has been exposed to experimental treatment or deliberately induced aging ^9,10^. This natural variation allows competence to be studied as an intrinsic property of the oocyte, rather than solely as a consequence of chronological aging or imposed stress. In goats, denuded oocytes lacking cumulus cells can still mature and give rise to embryos, but their developmental potential is lower than that of conventional cumulus-oocyte complexes and can be partly restored by co-culture with cumulus-enclosed oocytes^11^. These observations underscore the importance of the cumulus–oocyte interaction, but also indicate that the presence or absence of cumulus cells alone does not fully explain developmental competence. The cellular basis of the naturally occurring difference between morphologically excellent- and poor-quality oocytes therefore remains unclear.

The cumulus–oocyte complex provides an important metabolic and signalling environment during oocyte growth and maturation. Cumulus cells are highly glycolytic, whereas the oocyte relies substantially on oxidative metabolism, and metabolic communication between the two compartments influences meiotic and developmental competence^12^. Cumulus cells provide metabolic substrates and signalling molecules through gap-junctional communication and participate in the regulation of redox balance and meiotic progression^13^. Nevertheless, the oocyte must ultimately coordinate its own mitochondrial activity, energy production, redox state, calcium homeostasis and protein quality control during maturation^14,15^. Thus, naturally occurring differences in oocyte competence may reflect differences not only in somatic support but also in the metabolic and molecular state established within the oocyte itself. Identifying such intrinsic differences could help explain why some oocytes respond successfully to an otherwise supportive maturation environment whereas others do not.

Environmental stress provides an additional context in which oocyte quality and developmental competence can be compromised. Seasonal and thermal effects on reproductive outcomes have been reported across livestock species, with naturally occurring summer heat stress in cattle associated with a reduction in high-quality oocytes and an increase in low-quality oocytes ^16,17^. Importantly, emerging human evidence suggests that environmental temperature may also influence oocyte-related outcomes. Higher ambient temperatures before oocyte retrieval were recently associated with reduced numbers of total, mature and 2PN oocytes in women undergoing IVF/ICSI, while studies comparing seasonal IVF outcomes have reported changes in oocyte or embryo quality, although associations with cumulative live birth remain inconsistent ^18,19^. These observations suggest that the developing oocyte is sensitive to environmental conditions across species. However, the intracellular mechanisms linking cellular stress to the establishment of poor developmental competence remain poorly understood.

This question has often been approached through models of reproductive aging, in which oocyte quality progressively declines with maternal age. Such models have provided important insights into mitochondrial dysfunction, oxidative stress, spindle abnormalities and loss of developmental competence. They also implicate nicotinamide adenine dinucleotide (NAD⁺), a central metabolic cofactor involved in redox reactions, mitochondrial function, DNA repair and cellular stress responses. NAD⁺ availability influences the activity of NAD⁺-dependent enzymes, including sirtuins and poly (ADP-ribose) polymerases (PARPs), thereby linking cellular metabolism with mitochondrial function and chromatin regulation. Studies in aged mice have shown that NAD⁺ repletion, including supplementation with nicotinamide mononucleotide (NMN), can improve oocyte quality, mitochondrial function, meiotic competence and female fertility^20–22^. These findings establish a relationship between NAD⁺ metabolism and reproductive competence, but they do not address whether differences in NAD⁺ homeostasis are already present among otherwise contemporaneous oocytes before overt reproductive aging has occurred.

The distinction between naturally heterogeneous oocytes and aging oocytes is important because developmental competence may be influenced by metabolic state long before a measurable decline in ovarian function becomes apparent. An oocyte that is morphologically poor at collection may already have altered metabolic, organelle or stress-response pathways, but the nature and origin of such differences are not known. In particular, it is unclear whether NAD⁺ metabolism differs between naturally excellent- and poor-grade oocytes and, if so, whether such differences reflect altered NAD⁺ synthesis, increased consumption, or changes in the balance between these processes. Addressing NAD⁺ metabolism in this setting could therefore reveal cellular features associated with competence that are not readily apparent from morphology alone.

The de novo NAD⁺ biosynthetic pathway is of particular interest. NAD⁺ can be produced through several metabolic routes, including salvage and precursor-dependent pathways, while de novo synthesis converts tryptophan through the kynurenine pathway to quinolinic acid, which is subsequently converted to nicotinic acid mononucleotide (NAMN) by quinolinate phosphoribosyltransferase (QPRT). QPRT occupies a key position between quinolinate metabolism and downstream NAD⁺ synthesis^23,24^. Although reproductive studies have largely focused on NAD⁺ supplementation and salvage pathways, genetic studies have established that de novo NAD⁺ synthesis is relevant to ovarian function. Deletion of Qprt or Ido1 in mice reduces ovarian NAD⁺, impairs oocyte quality and fertility, and produces mitochondrial and spindle abnormalities; these phenotypes can be improved by NAD⁺ precursor supplementation^25^. These findings raise a more specific question for naturally heterogeneous oocytes: is de novo NAD⁺ biosynthesis already different between excellent- and poor-grade oocytes, and could a defect within this pathway contribute to the establishment of poor competence?

Cellular stress may provide an upstream context in which such metabolic differences arise. The endoplasmic reticulum (ER) coordinates protein folding, membrane homeostasis and intracellular calcium storage, and its function is closely integrated with mitochondrial activity^26^. Oocytes undergo extensive organelle remodeling and proteostatic changes during maturation, making maintenance of ER homeostasis particularly important^27^. When ER function is disturbed, the unfolded protein response (UPR), mediated principally by PERK, ATF6 and IRE1α, is activated to restore cellular homeostasis^28^. IRE1α is distinctive among the UPR sensors because, in addition to splicing XBP1 mRNA, its endoribonuclease can selectively cleave cellular transcripts through regulated IRE1-dependent decay (RIDD)^29,30^. This provides a potential mechanism by which ER stress could influence the abundance of metabolic enzymes and thereby alter cellular metabolism.

A connection between ER stress and de novo NAD⁺ synthesis has been described outside the reproductive system. In the kidney, cellular stress is associated with impaired QPRT-dependent NAD⁺ biosynthesis and quinolinate accumulation, and experimental evidence indicates that the ER-stress response can suppress QPRT expression^31^. Whether this relationship exists in the oocyte is unknown. More broadly, it is not clear whether the metabolic differences between naturally excellent- and poor-grade oocytes are accompanied by differences in ER homeostasis, whether ER stress is associated with altered de novo NAD⁺ biosynthesis, or whether IRE1α- dependent RNA regulation contributes to changes in metabolic competence. Finally, NAD⁺ metabolism provides a plausible point of convergence between several processes required for successful oocyte maturation. Changes in NAD⁺ availability could influence mitochondrial redox metabolism and ROS production, while mitochondrial dysfunction can in turn alter calcium handling and membrane potential. NAD⁺-dependent sirtuins and PARPs also provide direct links between metabolism, DNA repair and chromatin regulation. Thus, differences in NAD⁺ homeostasis could potentially be associated with coordinated changes in mitochondrial function, calcium balance, oxidative stress and chromatin state rather than with a single isolated phenotype. Whether this relationship holds in naturally poor-quality oocytes, and whether the resulting cellular state can be reversed, remains to be explored.

## Results

### Poor-grade oocytes exhibit an intrinsic defect in meiotic competence that is not rescued by cumulus-cell support

Oocyte morphology is routinely used as a practical indicator of oocyte quality; however, whether morphological abnormalities primarily reflect compromised somatic support or an intrinsic defect within the oocyte remains poorly understood. To distinguish between these possibilities, we first classified goat cumulus–oocyte complexes (COCs) according to established morphological criteria and subsequently compared their meiotic competence during in vitro maturation (IVM) (Fig. 1A & Fig. S1A).

**Figure 1.**
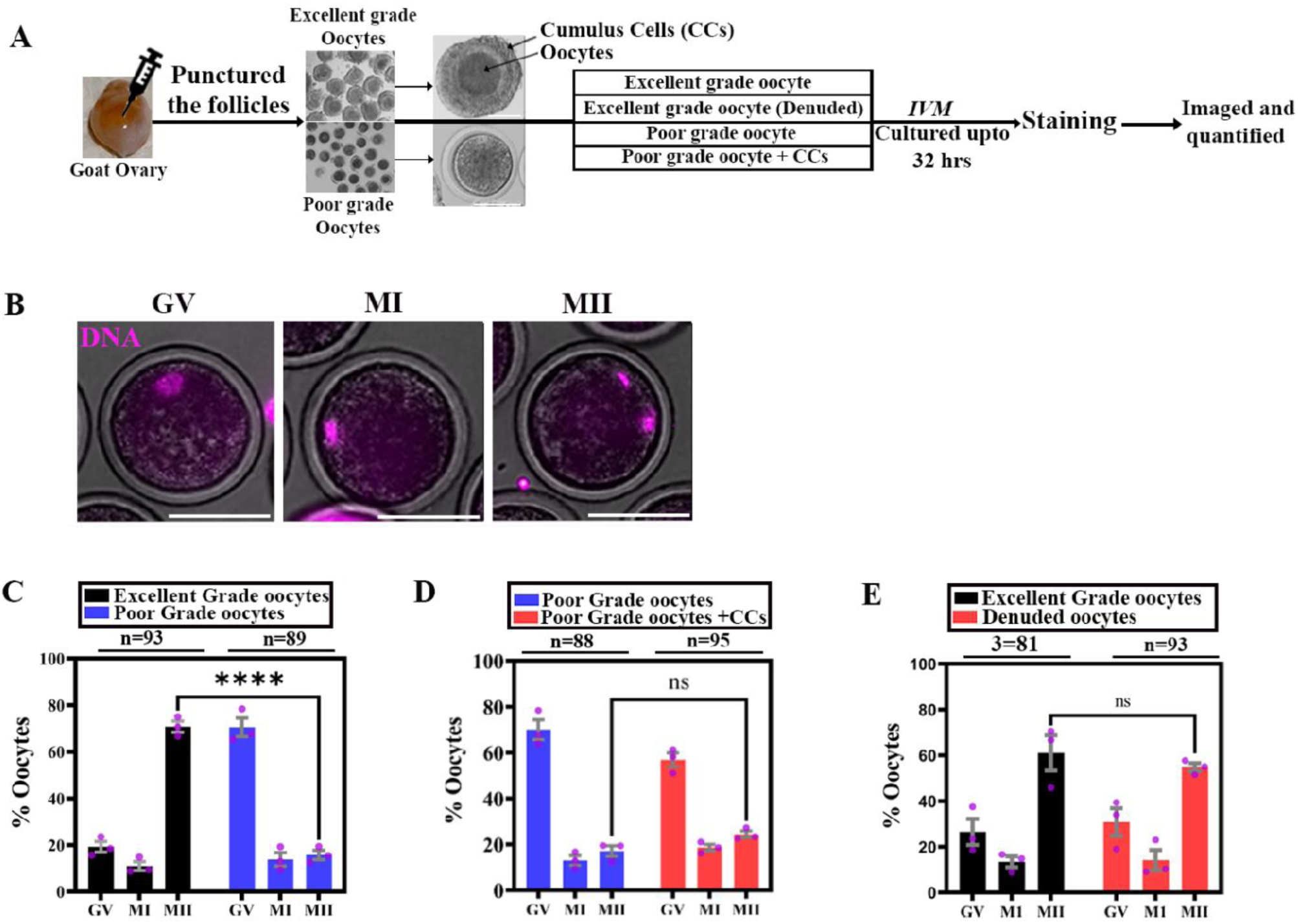
Poor-grade goat oocytes exhibit impaired meiotic competence independent of cumulus cell support. **(A)** Schematic of the experimental workflow. Oocytes were recovered from goat ovarian follicles, classified as excellent-grade or poor-grade based on morphology, and allocated to four groups: intact excellent-grade oocytes, denuded excellent-grade oocytes, poor-grade oocytes, and poor-grade oocytes co-cultured with cumulus cells. Oocytes were subjected to in vitro maturation (IVM) for up to 32 h, followed by staining and imaging. **(B)** Representative images of goat oocytes at the germinal vesicle (GV), metaphase I (MI), and metaphase II (MII) stages following DNA staining with DAPI (magenta). **(C)** Quantification of meiotic progression in excellent-grade (n = 93) and poor-grade (n = 89) oocytes, showing the proportions of oocytes classified as GV, MI, or MII after IVM. **(D)** Quantification of meiotic progression in poor-grade oocytes cultured without (n = 88) or with additional cumulus cells (n = 95). **(E)** Quantification of meiotic progression in intact excellent-grade (n = 81) and denuded excellent-grade (n = 93) oocytes. N denotes the number of independent biological replicates or experiments, and n denotes the total number of oocytes analyzed. Error bars show mean ± s.d., s.e.m. (n.s. = non-significant) (****) P ≤ 0.0001, Statistical significance was assessed using multiple unpaired t-test comparisons. Scale bars: 100 μm.

Excellent-grade oocytes were identified by compact and multilayered cumulus investment, homogeneous/granular cytoplasm, intact zona pellucida, and absence of visible cytoplasmic abnormalities, whereas poor-grade oocytes were characterized by sparse or no cumulus layers, and heterogeneous or dark/granular cytoplasm. Representative images of the two morphological classes are shown in Fig. 1A & Fig. S1A. Importantly, this classification was performed prior to IVM, allowing us to determine whether the morphological phenotype was predictive of subsequent meiotic competence.

We first asked whether differences in oocyte quality were associated with an intrinsic defect in meiotic progression. Following IVM, nuclear maturation was assessed by DAPI staining, and oocytes were classified as germinal vesicle (GV), metaphase I (MI), or metaphase II (MII) according to chromatin configuration (Fig. 1B). Excellent-grade oocytes exhibited robust meiotic progression, with approximately 70.75 ± 2.41% (s.e.m.) reaching the MII stage. In contrast, only 15.74 ± 1.99% (s.e.m.) of poor-grade oocytes progressed to MII, representing an approximately 4-fold reduction in maturation efficiency relative to excellent-grade oocytes (Fig. 1C). Because cumulus cells provide essential metabolic and paracrine support to the oocyte, we next asked whether the impaired maturation of poor-grade oocytes could be explained by inadequate somatic-cell support. Poor-grade oocytes were therefore co-cultured with cumulus cells throughout IVM (Fig. 1A). Additional cumulus-cell support failed to substantially restore meiotic competence, with 24.37 ± 1.46% (s.e.m.) of poor-grade oocytes reaching MII compared with 17.09 ± 2.30 (s.e.m.) in the corresponding control group (Fig. 1D & Fig. S1B). Thus, increasing the availability of cumulus-cell support was insufficient to overcome the maturation defect associated with poor oocyte morphology.

We then performed the reciprocal experiment using excellent-grade oocytes. Cumulus cells were mechanically removed before IVM to determine whether loss of somatic support was sufficient to impair maturation. Denuded excellent-grade oocytes retained high meiotic competence, with 54.73 ± 1.79% (s.e.m.) reaching MII compared with 61.02 ± 7.68% (s.e.m.) of cumulus-enclosed excellent-grade oocytes (Fig. 1E & Fig. S1C). The absence of a substantial reduction following cumulus removal further argues against inadequate cumulus- cell support as the primary determinant of the poor-grade phenotype. Together, these reciprocal experiments demonstrate that the reduced meiotic competence of poor-grade oocytes is unlikely to result solely from insufficient cumulus-cell support. Rather, poor morphological quality is associated with an oocyte-intrinsic defect in meiotic progression that is not readily rescued by restoring the surrounding somatic environment. These findings establish a functional distinction between excellent- and poor-grade oocytes and provide the rationale for investigating the intracellular molecular and metabolic alterations that underlie the loss of oocyte competence.

### Poor-grade oocytes exhibit a distinct metabolic state characterized by disruption of NAD⁺ metabolism

Having established in Figure 1 that poor-grade oocytes exhibit an intrinsic defect in meiotic competence that is not rescued by additional cumulus-cell support, we next sought to determine whether this functional difference is accompanied by an altered intracellular metabolic state. Because the oocytes were profiled at the GV stage, before meiotic maturation, we reasoned that metabolic differences detected at this stage could represent intrinsic features of poor-quality oocytes rather than secondary consequences of failed maturation. We therefore performed untargeted metabolomic profiling of GV-stage excellent- and poor-grade goat oocytes (Fig. 2A).

**Figure 2.**
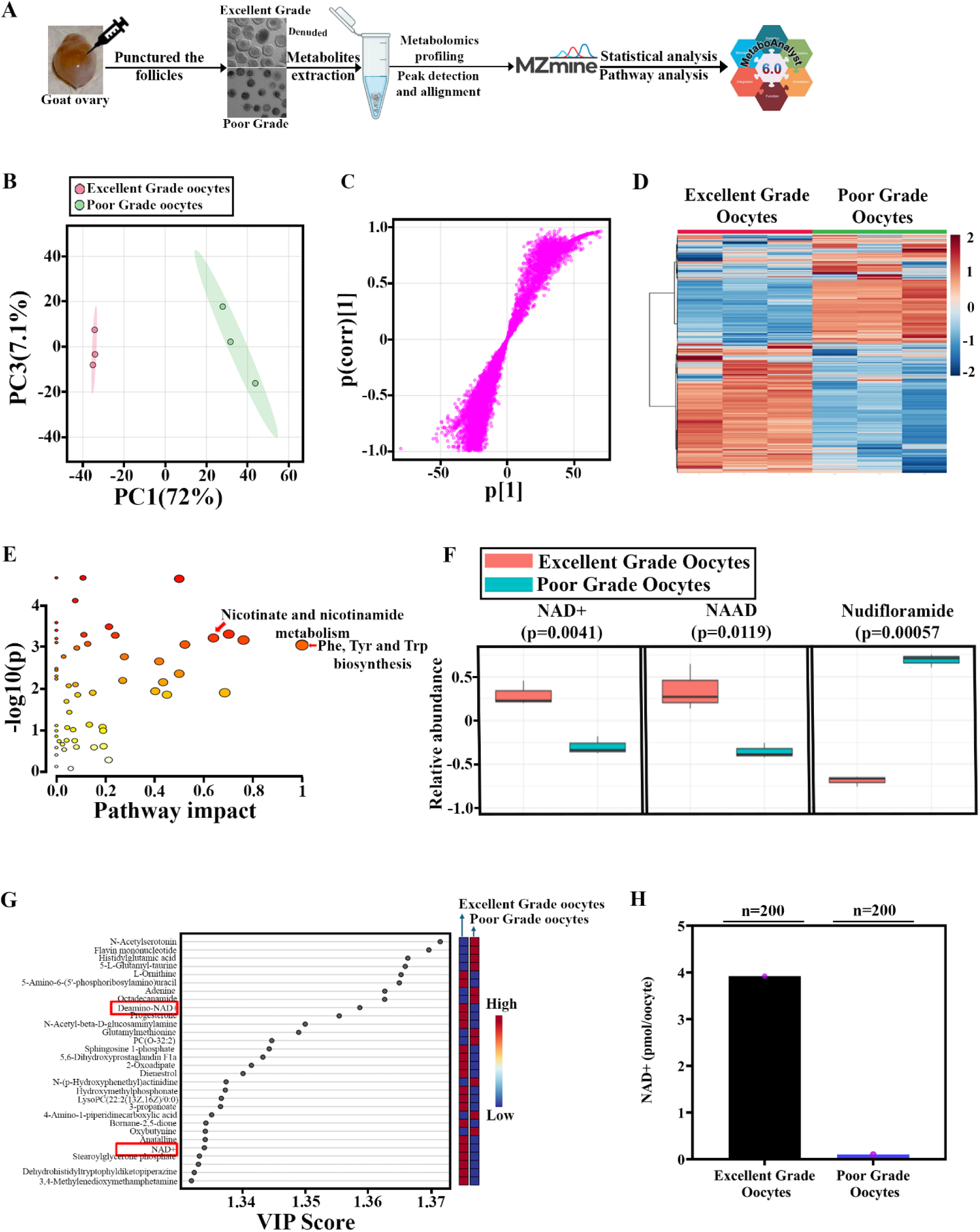
Untargeted metabolomic profiling reveals altered NAD⁺ metabolism in poor-grade goat oocytes. **(A)** Schematic of the experimental workflow for metabolomic profiling of excellent-grade and poor-grade goat oocytes. Metabolites were extracted from each group and analyzed by mass spectrometry, followed by peak detection, alignment, statistical analysis, and pathway enrichment analysis (N = 3 independent biological replicates; n = 750 oocytes per group). **(B)** Principal component analysis (PCA) showing separation of the metabolomic profiles of excellent-grade and poor-grade oocytes. PC1 and PC3 explain 72% and 7.1% of the total variance, respectively. **(C)** S-plot showing the covariance and correlation of individual metabolites with the discrimination between the two oocyte groups. **(D)** Heatmap showing the relative abundance of differential metabolites in excellent-grade and poor-grade oocytes. Colors represent relative metabolite abundance, with red and blue indicating higher and lower abundance, respectively. **(E)** Metabolite set enrichment and pathway topology analysis identifying metabolic pathways altered between excellent-grade and poor-grade oocytes. The x-axis represents pathway impact, and the y-axis represents the negative logarithm of the enrichment P-value. **(F)** Comparison of normalized abundance values for NAD⁺, nicotinic acid adenine dinucleotide (NAAD), and nudifloramide between excellent-grade and poor-grade oocytes, as determined by metabolomic analysis. P values are indicated above each comparison. **(G)** Variable importance in projection (VIP) scores for the top 50 metabolites contributing to discrimination between excellent-grade and poor-grade oocytes. The accompanying heatmap indicates relative metabolite abundance in each group. Metabolites with VIP scores greater than 1.3 are highlighted. **(H)** Quantification of NAD⁺ abundance in excellent-grade and poor-grade goat oocytes, showing reduced NAD⁺ levels in poor-grade oocytes (N = 1; n = 200 oocytes per group). N denotes the number of independent biological replicates or experiments, and n denotes the total number of oocytes analyzed. Statistical tests, data presentation, and multiple-testing correction are described in the Methods.

Following isolation from punctured follicles, excellent-grade oocytes were denuded, whereas poor-grade oocytes were analyzed in their isolated state, and metabolites were extracted for untargeted profiling. Unsupervised principal component analysis (PCA) revealed a clear separation between excellent- and poor-grade oocytes (Fig. 2B). PC1 accounted for 72% of the total variance, while PC3 accounted for 7.1%, with the two groups occupying distinct regions of the metabolic space. This separation indicates that oocyte quality is associated with a substantial difference in the intracellular metabolome rather than variation confined to a limited number of metabolites. Moreover, the close clustering of biological replicates within each group supports the reproducibility of the metabolic differences between excellent- and poor-grade oocytes. We next examined the contribution of individual metabolic features to the separation observed by PCA. The correlation/loading analysis demonstrated that numerous metabolic features contributed to the discrimination between the two oocyte populations (Fig. 2C), further supporting the presence of coordinated metabolic differences associated with oocyte quality rather than an effect driven by a single metabolite.

Consistent with the PCA, hierarchical clustering generated two distinct metabolic profiles corresponding to excellent- and poor-grade oocytes (Fig. 2D). The resulting heatmap revealed coordinated differences in metabolite abundance between the groups, with distinct clusters displaying opposing patterns across oocyte quality classes. Together, these complementary analyses demonstrate that poor-grade oocytes possess a distinct metabolic state that is already evident at the GV stage, before meiotic maturation. This finding prompted us to determine which metabolic pathways contribute most strongly to the metabolic divergence associated with poor oocyte quality.

To determine which metabolic pathways were most strongly associated with this separation, we performed pathway enrichment and impact analysis of the altered metabolites. Among the pathways identified, nicotinate and nicotinamide metabolism showed a prominent pathway impact and significant enrichment (Fig. 2E). Notably, the analysis also identified phenylalanine, tyrosine and tryptophan biosynthesis as an altered pathway. The convergence of these metabolic alterations was particularly notable because tryptophan metabolism provides an important route for de novo NAD⁺ biosynthesis, whereas nicotinate/nicotinamide metabolism encompasses multiple NAD⁺ precursor and biosynthetic reactions. This pathway-level convergence prompted us to examine individual NAD⁺-related metabolites. Three metabolites showed particularly notable differences between the two oocyte populations:

NAD⁺, NAAD (deamido-NAD), and nudifloramide (Fig. 2F). NAD⁺ and NAAD were significantly reduced in poor-grade oocytes, with P = 0.0041 and P = 0.0119, respectively, whereas nudifloramide was markedly increased in poor-grade oocytes (P = 0.00057). Thus, the metabolomic phenotype was not limited to a reduction in NAD⁺ itself but involved coordinated changes in multiple metabolites associated with NAD-related metabolism. The importance of the NAD⁺-associated metabolic signature was further supported by supervised variable-importance analysis. The VIP analysis identified NAAD/deamino-NAD and NAD⁺ among the discriminating metabolites contributing to the separation of excellent- and poor-grade oocytes (Fig. 2G). The convergence of pathway enrichment, metabolite-level analysis, and multivariate discrimination therefore independently pointed toward NAD⁺ metabolic homeostasis as a major metabolic feature associated with poor oocyte quality.

Because untargeted metabolomics provides relative rather than absolute metabolite abundance, we next independently quantified NAD⁺ levels in excellent- and poor-grade oocytes (Fig. 2H). Consistent with the metabolomic analysis, excellent-grade oocytes contained approximately 3.92 pmol NAD⁺/oocyte, whereas poor-grade oocytes contained only approximately 0.10 pmol NAD⁺/oocyte, corresponding to an approximately 39.2-fold (97.4%) reduction in NAD⁺ abundance in poor-grade oocytes (Fig. 2H). Thus, three complementary levels of analysis, global metabolic separation, pathway enrichment, and targeted measurement of NAD⁺ converged on the same metabolic phenotype. Poor-grade oocytes are therefore characterized by a profound alteration of intracellular metabolism, with NAD⁺ depletion emerging as one of the most prominent and reproducible metabolic abnormalities.

### Poor-grade oocytes exhibit a selective defect in de novo NAD⁺ biosynthesis

The marked reduction in NAD⁺ abundance in poor-grade oocytes raised the question of whether this metabolic deficiency resulted from excessive NAD⁺ consumption, insufficient biosynthesis, or a combination of both. Because NAD⁺ homeostasis is determined by the balance between its synthesis and utilization, we first examined whether limiting NAD⁺ consumption could improve the meiotic competence of poor-grade oocytes (Fig. 3A, B). Poor- grade oocytes were subjected to IVM in the presence or absence of quercetin, and meiotic progression was assessed by quantifying the proportion of oocytes reaching the MII stage. Quercetin treatment produced a significant but relatively modest increase in MII-stage oocytes, from approximately 15.9 ± 3.62% (s.e.m.) in untreated poor-grade oocytes to 33.7 ± 1.81% (s.e.m.) following treatment (P < 0.05; Fig. 3C, Fig. & Fig. S3A). Although this finding indicates that NAD⁺ availability contributes to meiotic progression, the limited extent of rescue suggested that increased NAD⁺ consumption was unlikely to be the predominant explanation for the profound NAD⁺ deficiency observed in poor-grade oocytes.

**Figure 3.**
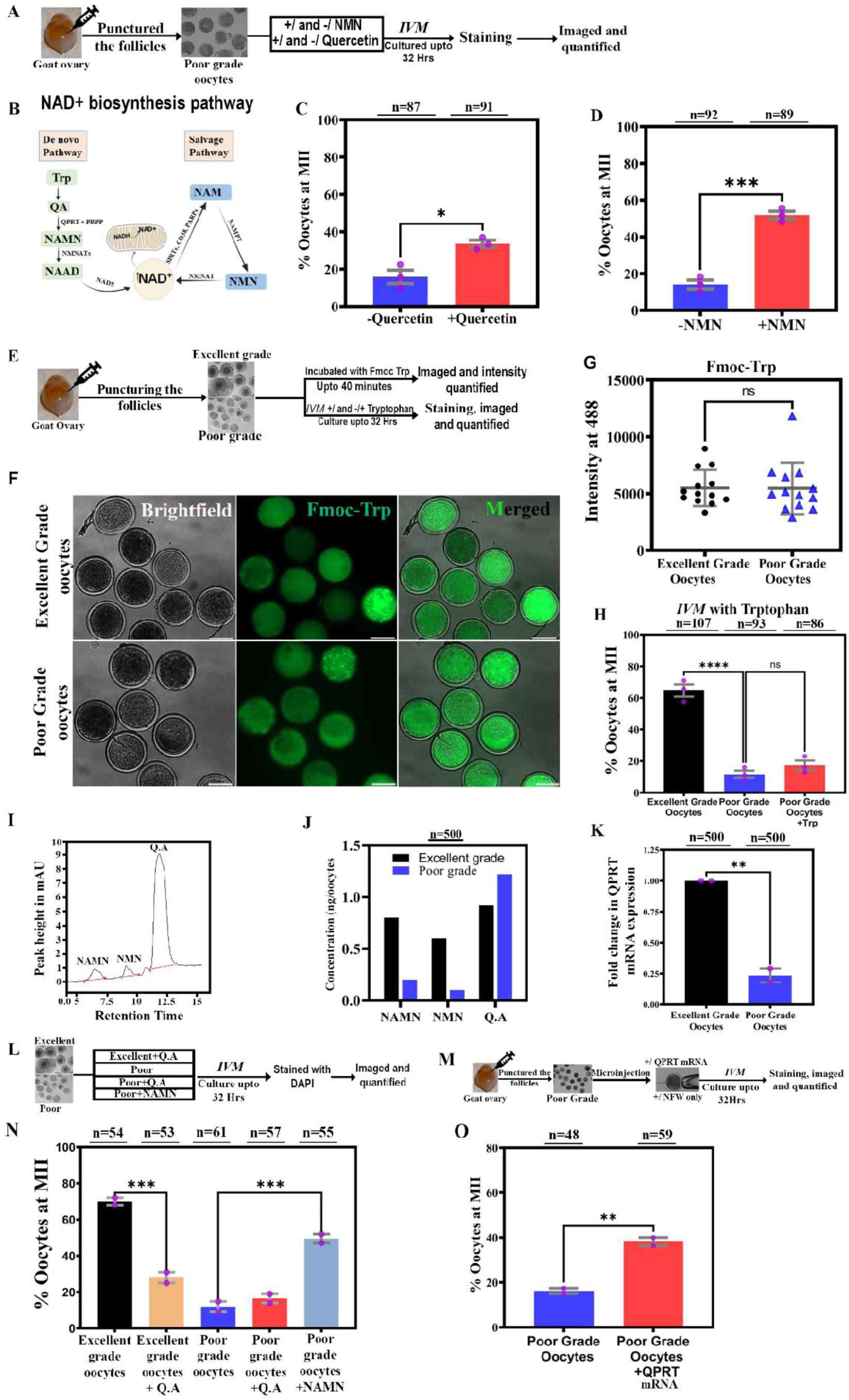
Functional analysis identifies impaired de novo NAD⁺ biosynthesis at the QPRT-dependent step in poor-grade goat oocytes. **(A)** Schematic of the experimental workflow for in vitro maturation (IVM) of poor-grade goat oocytes treated with NMN and/or quercetin, followed by staining and assessment of meiotic maturation. **(B)** Schematic representation of the de novo and salvage pathways of NAD⁺ biosynthesis. The de novo pathway converts tryptophan-derived intermediates into NAD⁺ through the quinolinate phosphoribosyltransferase (QPRT)-dependent step, whereas the salvage pathway regenerates NAD⁺ from nicotinamide through NMN. **(C)** Quantification of the proportion of poor-grade oocytes reaching metaphase II (MII) following IVM in the absence (n = 87) or presence (n = 91) of quercetin. **(D)** Quantification of MII-stage maturation in poor-grade oocytes cultured without (n = 92) or with (n = 89) NMN supplementation. **(E)** Schematic of the experimental workflow used to assess tryptophan uptake and meiotic maturation in excellent-grade and poor-grade goat oocytes. Oocytes were incubated with fluorescently labelled tryptophan (Fmoc-Trp), and intracellular fluorescence was quantified. Parallel groups were cultured with or without tryptophan supplementation during IVM. **(F)** Representative bright-field, Fmoc-Trp fluorescence, and merged images of excellent-grade and poor-grade goat oocytes following incubation with fluorescently labelled tryptophan. **(G)** Quantification of intracellular Fmoc-Trp fluorescence intensity in excellent-grade and poor-grade oocytes. No significant difference was detected between the groups. **(H)** Quantification of the proportion of excellent-grade and poor-grade oocytes reaching MII following IVM, with or without exogenous tryptophan supplementation. Tryptophan supplementation did not significantly improve maturation of poor-grade oocytes. **(I)** Representative HPLC chromatogram showing the retention profiles of quinolinic acid (QA), nicotinic acid mononucleotide (NAMN), and nicotinamide mononucleotide (NMN). **(J)** Comparison of NAMN, NMN, and QA abundance in excellent-grade and poor-grade oocytes, showing reduced NAMN and NMN and increased QA in poor-grade oocytes. **(K)** Relative QPRT mRNA expression in excellent-grade and poor-grade goat oocytes, assessed by RT-qPCR. **(L)** Schematic of the experimental design used to assess whether supplementation with QA or NAMN improves meiotic maturation in poor-grade oocytes. Maturation rates were compared among excellent-grade controls, poor-grade controls, and poor-grade oocytes supplemented with QA or NAMN. **(M)** Schematic of QPRT mRNA microinjection into poor-grade goat oocytes, followed by IVM and assessment of meiotic maturation. **(N)** Quantification of the proportion of oocytes reaching MII following treatment with QA or NAMN, compared with excellent-grade and untreated poor-grade controls. **(O)** Quantification of MII-stage maturation in poor-grade oocytes following microinjection of QPRT mRNA compared with untreated poor-grade controls. Data are presented as mean ± SEM, where applicable. The number of oocytes analyzed for each group is indicated above the corresponding bars. Statistical significance is indicated by asterisks (*P < 0.05, **P < 0.01, ***P < 0.001, ****P < 0.0001, n.s. = non-significant. Statistical significance was assessed using unpaired t-test and ordinary one-way ANOVA **(H and L).** Scale bars: 100 μ m Scale.

We therefore asked whether increasing NAD⁺ biosynthetic capacity would have a more pronounced effect. Because nicotinamide mononucleotide (NMN) can serve as a precursor for NAD⁺ biosynthesis through the salvage pathway (Fig. 3B), poor-grade oocytes were supplemented with NMN during IVM (Fig. 3A). In contrast to the modest effect of quercetin, NMN supplementation produced a notable improvement in meiotic maturation, increasing the proportion of MII-stage oocytes from 14.06 ± 2.36% (s.e.m.) to 51.83 ± 2.10% (s.e.m.) (P < 0.001; Fig. 3D & Fig. S3B). The substantially greater rescue following NMN supplementation suggested that inadequate NAD⁺ production, rather than excessive NAD⁺ consumption alone, may represent a major metabolic limitation in poor-grade oocytes.

We next sought to determine which branch of NAD⁺ biosynthesis was compromised. Mammalian cells can generate NAD⁺ through both salvage and de novo pathways, with tryptophan serving as the initiating substrate for the latter (Fig. 3B). We therefore first examined whether poor-grade oocytes had impaired access to tryptophan. Excellent- and poor-grade oocytes were incubated with fluorescently labeled Fmoc-tryptophan for up to 40 min, and intracellular fluorescence was visualized and quantified (Fig. 3E,F). Both excellent- and poor-grade oocytes displayed readily detectable intracellular Fmoc-tryptophan, and quantitative analysis revealed no significant difference in fluorescence intensity between the two groups (Intensity: 5498 ± 1595 (s.d.) vs 5467 ± 2280 (s.d.); Fig. 3G). Thus, the reduced NAD⁺ state of poor-grade oocytes was not attributable to an obvious defect in tryptophan uptake.

We next tested whether increasing the availability of tryptophan could overcome the impaired maturation phenotype. Supplementation with exogenous tryptophan during IVM, however, failed to improve meiotic progression of poor-grade oocytes (Fig. 3H). Whereas 64.73 ± 3.89% (s.e.m.) of excellent-grade oocytes reached MII, only 11.67 ± 2.33% (s.e.m.) of untreated poor-grade oocytes reached MII, and tryptophan supplementation resulted in approximately 17.33 ± 2.96% (s.e.m.) MII oocytes, with no significant difference between untreated and tryptophan-treated poor-grade oocytes (Fig. 3H). These findings indicated that tryptophan availability was unlikely to be the limiting factor and instead pointed to a defect downstream within the de novo NAD⁺ biosynthetic pathway.

To localize this defect further, we examined metabolites along the de novo pathway. Targeted analysis detected NAMN, NMN, and quinolinic acid (QA) in oocyte extracts (Fig. 3I & Fig. S3D). Notably, poor-grade oocytes exhibited a pronounced accumulation of QA accompanied by reduced levels of the downstream intermediates NAMN and NMN compared with excellent-grade oocytes (Fig. 3J & Fig. S3E). This reciprocal metabolic pattern was consistent with impaired utilization of QA for downstream NAD⁺ biosynthesis and suggested a bottleneck at the conversion of QA to NAMN. This reaction is catalyzed by quinolinate phosphoribosyltransferase (QPRT), prompting us to examine QPRT in poor-grade oocytes.

Consistent with the metabolite profile, QPRT expression was markedly reduced in poor-grade oocytes, reaching approximately 4.26-fold of the level detected in excellent-grade oocytes (P < 0.01; Fig. 3K). The combination of QA accumulation, depletion of downstream intermediates, and reduced QPRT expression therefore localized the metabolic abnormality to the QPRT-dependent step of the de novo NAD⁺ biosynthetic pathway. We next tested whether manipulating metabolites on either side of this putative metabolic bottleneck could influence meiotic competence. Addition of exogenous QA significantly reduced maturation of excellent-grade oocytes, decreasing the proportion reaching MII from approximately 70.0 ± 2.0% (s.e.m.) to 28.0 ± 3.0% (s.e.m.) (P < 0.001; Fig. 3L,M & Fig. S3F). In contrast, addition of QA to poor-grade oocytes did not further significantly alter their already compromised maturation, with approximately 11.91 ± 2.9% (s.e.m.) and 16.5 ± 2.5% (s.e.m.) of untreated and QA-treated poor-grade oocytes, respectively, reaching MII (Fig. 3L,M & Fig. S3F). This finding is consistent with poor-grade oocytes already possessing elevated endogenous QA. Importantly, supplementation with NAMN, the metabolic product immediately downstream of the QPRT reaction, substantially improved maturation of poor-grade oocytes, increasing MII progression from approximately 11.91 ±% (s.e.m.) to 49.5 ± 2.5% (s.e.m.) (P < 0.001; Fig. 3L,M & Fig. S3F). Thus, bypassing the putative QPRT-dependent metabolic bottleneck was sufficient to restore a substantial proportion of the meiotic competence lost in poor-grade oocytes.

Finally, to directly test whether reduced QPRT contributes to the maturation defect, we restored QPRT expression by microinjecting QPRT mRNA into poor-grade oocytes before IVM (Fig. 3N). QPRT restoration significantly increased the proportion of poor-grade oocytes reaching MII, from approximately 16.20 ± 1.95% (s.e.m.) in untreated oocytes to 38.18 ± 1.82% (s.e.m.) following QPRT mRNA supplementation (P < 0.01; Fig. 3O & Fig. S3G). This genetic rescue strengthens the evidence that reduced QPRT availability is functionally linked to the metabolic and meiotic defects of poor-grade oocytes. Taken together, these findings progressively establish QPRT-dependent de novo NAD⁺ biosynthesis as a critical metabolic determinant of poor-grade oocyte competence and provide a mechanistic link between the NAD⁺ deficiency and the intrinsic meiotic defect.

### ER stress and IRE1α signalling are associated with reduced QPRT expression and impaired meiotic competence in poor-grade oocytes

The accumulation of quinolinic acid, reduction of downstream NAD⁺ biosynthetic intermediates and decreased QPRT expression in poor-grade oocytes suggested a defect in the regulation of de novo NAD⁺ biosynthesis. To investigate the cellular processes associated with this phenotype, we performed comparative LC–MS/MS proteomic analysis of denuded excellent-grade and poor-grade goat oocytes collected from punctured follicles (Fig. 4A). Of the proteins identified, 17,566 were shared between the two groups, whereas 2617 and 2509 proteins were detected exclusively in excellent-grade and poor-grade oocytes, respectively (Fig. 4B). Principal component analysis separated the two groups, with PC1 and PC2 accounting for 45.2% and 21.0% of the total variance, respectively (Fig. 4C). A complementary component analysis also distinguished the two populations, with components 1 and 2 accounting for 21.9% and 36.9% of the variance, respectively (Fig. 4D). Hierarchical clustering further revealed distinct protein-abundance patterns between excellent-grade and poor-grade oocytes (Fig. 4E), indicating that differences in oocyte quality are accompanied by broad proteomic changes.

**Figure 4.**
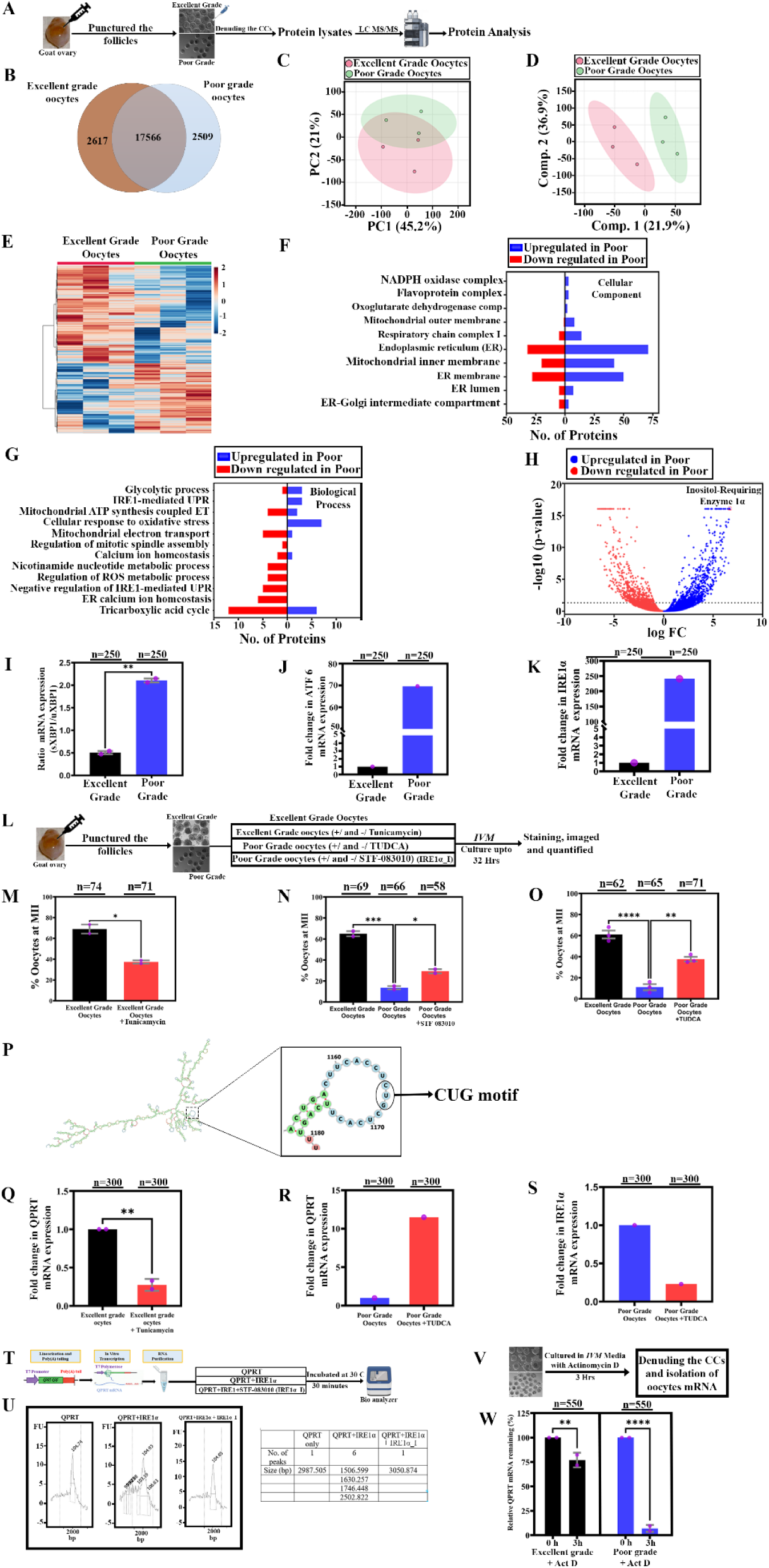
Proteomic profiling identifies ER stress-associated signaling and links IRE1α activation to QPRT dysregulation in poor-grade goat oocytes. **(A)** Schematic of the workflow for comparative proteomic analysis of denuded excellent-grade and poor-grade goat oocytes, including protein extraction, LC–MS/MS analysis, and downstream protein profiling. **(B)** Venn diagram showing the overlap between proteins identified in excellent-grade and poor-grade oocytes. **(C)** Principal component analysis (PCA) showing separation of the proteomic profiles of excellent-grade and poor-grade oocytes. **(D)** Partial least squares discriminant analysis (PLS-DA) score plot illustrating differences in protein profiles between the two groups. **(E)** Heatmap showing the relative abundance of differentially abundant proteins in excellent-grade and poor-grade oocytes. **(F,G)** Gene Ontology (GO) enrichment analysis of differentially abundant proteins associated with cellular components (F) and biological processes (G). Blue bars indicate proteins upregulated in poor-grade oocytes, whereas red bars indicate proteins downregulated in poor-grade oocytes. **(H)** Volcano plot showing differential protein abundance between excellent-grade and poor-grade oocytes, highlighting significantly altered proteins, including IRE1α. **(I–K)** RT-qPCR analysis of ER stress-associated transcripts in excellent-grade and poor-grade oocytes: (I) the ratio of spliced to unspliced XBP1 mRNA (XBP1s/XBP1u), (J) relative ATF6 mRNA expression, and (K) relative IRE1α (ERN1) mRNA expression. Each group comprised 250 oocytes. **(L)** Schematic of the experimental design used to assess meiotic maturation following treatment with tunicamycin ™ in excellent-grade oocytes, or tauroursodeoxycholic acid (TUDCA) and the IRE1α inhibitor STF-083010 (STF) in poor-grade oocytes, during in vitro maturation (IVM). **(M–O)** Quantification of the proportion of oocytes reaching metaphase II (MII) following pharmacological manipulation of ER stress signaling: (M) excellent-grade oocytes treated with or without TM; (N) poor-grade oocytes treated with or without STF; and (O) poor-grade oocytes treated with or without TUDCA. The numbers of oocytes analyzed are indicated above the bars. **(P)** Predicted secondary structure of QPRT mRNA showing a candidate stem– loop region containing a CUG motif. This computational prediction identifies a potential RNA structural feature for further investigation. **(Q)** Relative QPRT mRNA expression in excellent-grade oocytes and excellent-grade oocytes treated with TM, assessed by RT-qPCR. **®** Relative QPRT mRNA expression in poor-grade oocytes following TUDCA treatment. **(S)** Relative IRE1α (*ERN1*) mRNA expression in poor-grade oocytes following TUDCA treatment. **(T)** Schematic of the in vitro transcription and preparation of QPRT RNA substrates for the IRE1α-mediated RNA cleavage assay. **(U)** Representative RNA cleavage assay showing QPRT RNA incubated under the indicated conditions, including recombinant IRE1α. The accompanying table lists the sizes of the detected RNA products. **(V)** Schematic of the experimental workflow for assessing QPRT mRNA stability in excellent-grade and poor-grade oocytes following actinomycin D treatment to inhibit new transcription. **(W)** Relative QPRT mRNA abundance in excellent-grade and poor-grade oocytes at 0 and 3 h following actinomycin D treatment, assessed by RT-qPCR. Values are expressed relative to the corresponding 0-hr level. The number of oocytes analyzed is indicated above each graph. Data are presented as mean ± SEM. Statistical significance is indicated by asterisks (*P < 0.05, **P < 0.01, ***P < 0.001, ****P < 0.0001. Statistical significance was assessed using unpaired t-test and ordinary one-way ANOVA **(N and O)** and 2way ANOVA **(W).**

Functional annotation of the differentially represented proteins identified cellular components associated with the endoplasmic reticulum (ER), including the ER membrane and lumen, as well as the mitochondrial inner membrane, respiratory-chain complex I and NADPH oxidase complex (Fig. 4F). Biological-process analysis highlighted changes in pathways associated with the IRE1-mediated unfolded protein response, cellular responses to oxidative stress, mitochondrial ATP synthesis and electron transport, calcium homeostasis, nicotinamide nucleotide metabolism, ROS regulation and the tricarboxylic acid cycle (Fig. 4G). These findings indicated that poor-grade oocytes exhibit coordinated changes in ER-associated, mitochondrial and metabolic processes. The enrichment of IRE1-mediated unfolded protein response pathways prompted us to examine whether IRE1α signalling was altered in poor-grade oocytes.

We first quantified the ratio of spliced to unspliced XBP1 mRNA as a readout of IRE1α-dependent RNA processing. The XBP1s/XBP1u ratio increased from 0.5 in excellent-grade oocytes to 2.1 in poor-grade oocytes, corresponding to an approximately 4.17-fold increase (Fig. 4I). We next examined the expression of ATF6 and IRE1α transcripts. ATF6 mRNA showed an approximately 70-fold increase, while IRE1α mRNA increased approximately 240-fold in poor-grade oocytes relative to excellent-grade controls (Fig. 4J,K). The increased XBP1 splicing, together with the transcript-expression results and proteomic pathway analysis, supports the involvement of ER-stress signalling, particularly the IRE1α–XBP1 axis, in poor-grade oocytes. However, changes in transcript abundance alone do not establish activation of the corresponding proteins.

To determine whether ER stress could influence meiotic competence, we exposed excellent-grade oocytes to tunicamycin during in vitro maturation (IVM), while poor-grade oocytes were treated with either TUDCA, a chemical chaperone used to alleviate ER stress, or STF-083010, an inhibitor of IRE1α signalling (Fig. 4L). Tunicamycin reduced the proportion of excellent-grade oocytes reaching metaphase II (MII) from 68.98 ± 4.34% (s.e.m.) to 37.35 ± 1.54% (s.e.m.) (Fig. 4M & Fig. S4A). Conversely, treatment with STF-083010 increased MII progression in poor-grade oocytes from 13.58 ± 1.58% (s.e.m.) to 29.3 ± 1.95% (s.e.m.) (Fig. 4N). TUDCA treatment similarly improved maturation in poor-grade oocytes, increasing the proportion reaching MII from 11.33 ± 2.89% (s.e.m.) to 37.67 ± 2.19% (s.e.m.) (Fig. 4O & Fig. S4B). These results indicate that experimentally induced ER stress can impair meiotic maturation and that pharmacological attenuation of ER stress or IRE1α signalling can improve the meiotic competence of poor-grade oocytes. The effects of these interventions support a functional contribution of ER-stress-associated processes to the poor-grade phenotype, although they do not establish that IRE1α signalling accounts for all the effects of ER stress. Because IRE1α possesses endoribonuclease activity and can promote regulated IRE1-dependent decay (RIDD) of selected transcripts, we investigated whether QPRT RNA might be susceptible to IRE1α-mediated processing. Secondary-structure prediction identified a structured region containing a CUG motif in the QPRT transcript (Fig. 4P). This provided a candidate region for investigating potential RNA recognition, although the predicted structure alone does not establish direct recognition by IRE1α.

We next examined QPRT expression following ER-stress manipulation. Tunicamycin treatment reduced QPRT mRNA abundance in excellent-grade oocytes to 3.7-fold relative to the untreated reference (Fig. 4Q). Conversely, TUDCA treatment increased QPRT mRNA abundance in poor-grade oocytes 11-fold relative to untreated poor-grade controls (Fig. 4R). IRE1α mRNA abundance decreased following TUDCA treatment to 4.35-fold (Fig. 4S). These results link ER-stress manipulation with changes in QPRT expression, although they do not distinguish direct RNA cleavage from indirect transcriptional or post-transcriptional regulation.

To determine whether IRE1α can directly process QPRT RNA, we incubated an in-vitro-transcribed QPRT RNA substrate with recombinant IRE1α in the presence or absence of STF-083010 (Fig. 4T). RNA fragment profiles were analysed using an Agilent 2100 Bioanalyzer.

In the absence of recombinant IRE1α, the profile was dominated by the intact QPRT RNA product, whereas incubation with IRE1α resulted in additional RNA products consistent with RNA cleavage (Fig. 4U & Fig. S4CD). Together, these findings indicate that recombinant IRE1α can process QPRT RNA in vitro and that this processing is sensitive to STF-083010. However, whether IRE1α mediates QPRT RNA cleavage in intact oocytes remains to be established. Finally, we examined QPRT RNA stability using an actinomycin D chase experiment (Fig. 4V). QPRT mRNA abundance declined following transcriptional inhibition in both excellent-grade and poor-grade oocytes, but the reduction was more pronounced in poor-grade oocytes after 3 h. Relative QPRT mRNA abundance decreased to 77% in excellent-grade oocytes and to 7% in poor-grade oocytes (Fig. 4W). These results indicate that QPRT RNA is less stable in poor-grade oocytes under the conditions examined. Collectively, the proteomic and functional analyses identify ER-stress-associated signalling as a potential contributor to the reduced QPRT expression observed in poor-grade oocytes. Increased XBP1 splicing, the effects of pharmacological ER-stress modulation on meiotic maturation, changes in QPRT expression following treatment, and the in-vitro RNA-processing results support a possible connection between IRE1α signalling and QPRT RNA stability. In the context of the observed accumulation of QA and depletion of downstream NAD⁺ biosynthetic intermediates, these findings support a model in which ER-stress-associated regulation of QPRT may contribute to impaired de novo NAD⁺ biosynthesis and reduced meiotic competence.

### NAD⁺ restoration alleviates calcium dysregulation and mitochondrial dysfunction, improves spindle morphology-associated outcomes, and supports early embryonic development in poor-grade oocytes

The reduced NAD⁺ levels and altered ER stress signalling observed in poor-grade oocytes prompted us to investigate whether these changes were accompanied by broader defects in cellular homeostasis. The endoplasmic reticulum (ER) regulates intracellular calcium storage, while mitochondria depend on tightly controlled calcium handling and redox balance to maintain their function. Because these processes are also important for meiotic spindle organization and oocyte developmental competence, we examined ER-associated fluorescence, intracellular calcium, oxidative stress, mitochondrial status and spindle morphology in excellent- and poor-grade goat oocytes.

Denuded oocytes were stained with complementary fluorescent probes to assess these cellular features (Fig. 5A). ER-Tracker staining revealed higher fluorescence intensity in poor-grade oocytes than in excellent-grade oocytes (Intensity: 1312 ± 345.6 (s.d.) vs 1032 ± 152.6 (s.d.); poor vs excellent; Fig. 5B,C), indicating an alteration in ER-associated fluorescence. We next assessed cytosolic calcium using Fluo-3 AM. Fluorescence intensity was higher in poor-grade oocytes than in excellent-grade oocytes (Intensity: 705 ± 114.2 (s.d.) vs 597 ± 146.6 (s.d.); poor vs excellent; Fig. 5F,G), consistent with increased cytosolic Ca²⁺ levels. A similar difference was observed with Rhod-2 AM, which showed higher fluorescence in poor-grade oocytes (Intensity: 2706 ± 595 (s.d.) 2223 ± 425 (s.d.); poor vs excellent; Fig. 5H,I), suggesting increased mitochondrial Ca²⁺ accumulation. These findings indicate that poor-grade oocytes exhibit altered calcium homeostasis in both the cytosolic and mitochondrial compartments.

**Figure 5.**
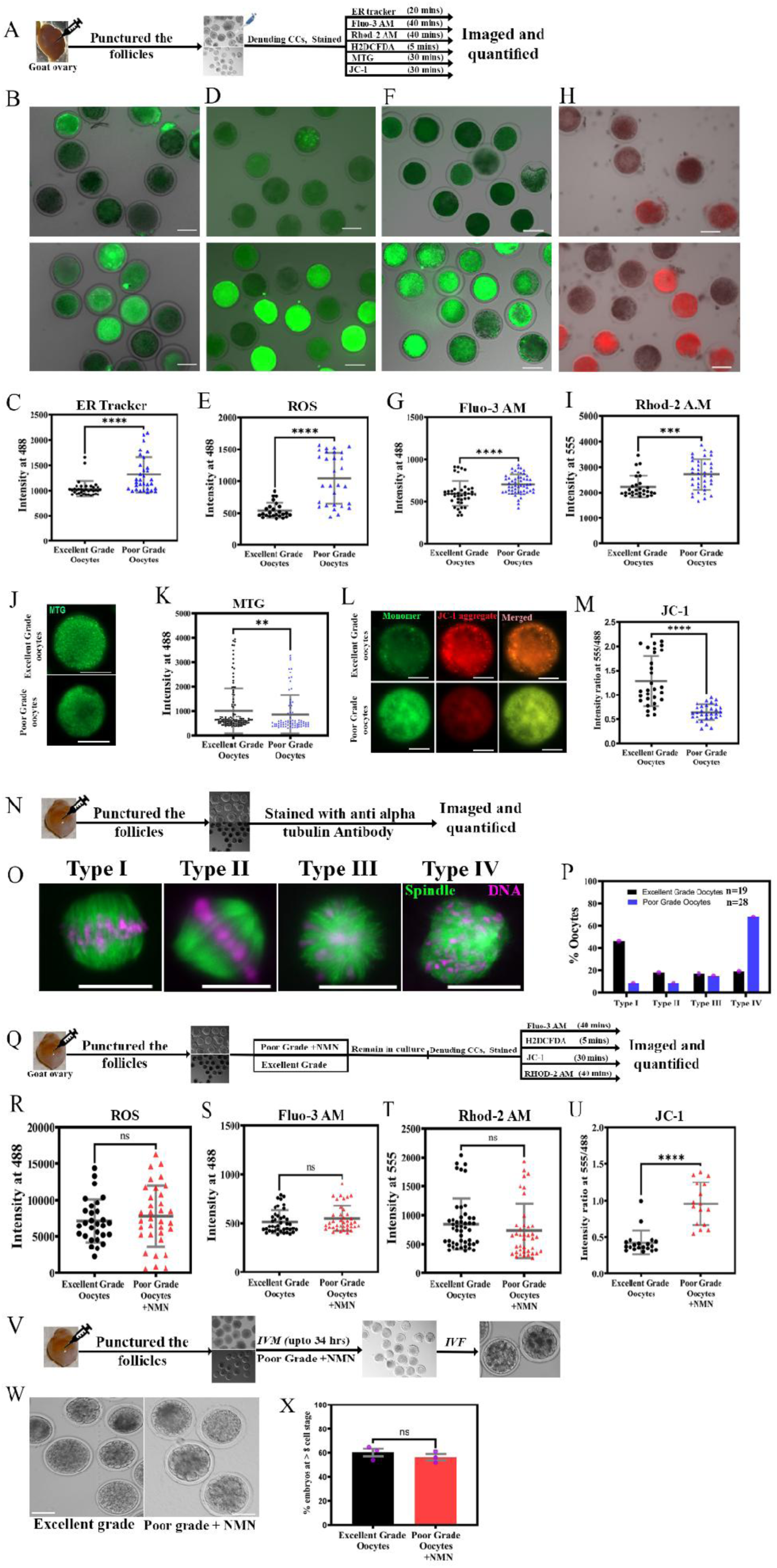
Poor-grade goat oocytes exhibit ER-associated calcium dysregulation, oxidative stress, and mitochondrial dysfunction, which are partially alleviated by NMN supplementation. **(A)** Schematic of the experimental workflow for isolating goat oocytes, performing fluorescence staining, and acquiring and quantifying fluorescence images. **(B, C)** Representative fluorescence images (B) and quantification of ER-Tracker fluorescence intensity (C) in excellent-grade and poor-grade oocytes, showing altered ER staining patterns and increased fluorescence intensity in poor-grade oocytes. **(D, E)** Representative fluorescence images (D) and quantification of intracellular reactive oxygen species (ROS) using H₂DCFDA staining (E) in excellent-grade and poor-grade oocytes. **(F, G)** Representative fluorescence images (F) and quantification of intracellular Ca²⁺ levels using Fluo-3 AM staining (G) in excellent-grade and poor-grade oocytes. **(H, I)** Representative fluorescence images (H) and quantification of mitochondrial Ca²⁺ levels using Rhod-2 AM staining (I) in excellent-grade and poor-grade oocytes. **(J, K)** Representative MitoTracker Green (MTG) fluorescence images (J) and quantification of fluorescence intensity (K) in excellent-grade and poor-grade oocytes, assessing mitochondrial distribution and signal intensity. **(L, M)** Representative JC-1 fluorescence images showing the green monomer and red aggregate signals, together with the merged images (L), and quantification of the red-to-green fluorescence ratio (M) in excellent-grade and poor-grade oocytes. A lower ratio indicates reduced mitochondrial membrane potential. **(N)** Schematic of the experimental workflow for assessing meiotic spindle morphology in goat oocytes using immunofluorescence staining for α-tubulin and DNA staining. **(O)** Representative images of the four meiotic spindle morphology categories (Types I–IV). Spindle (green) and DNA (magenta). **(P)** Quantification of the distribution of spindle morphology categories (Types I–IV) in excellent-grade and poor-grade oocytes. **(Q)** Schematic of the experimental workflow for evaluating the effects of NMN supplementation on ROS levels, intracellular Ca²⁺, mitochondrial Ca²⁺, and mitochondrial membrane potential in poor-grade oocytes, with excellent-grade oocytes serving as controls. **(R–U)** Quantification of intracellular ROS levels using H₂DCFDA (R), intracellular Ca²⁺ levels using Fluo-3 AM (S), mitochondrial Ca²⁺ levels using Rhod-2 AM (T), and mitochondrial membrane potential using the JC-1 red-to-green fluorescence ratio (U) in excellent-grade oocytes and NMN-treated poor-grade oocytes. **(V)** Schematic of the experimental workflow used to assess subsequent embryonic development following IVM of untreated poor-grade oocytes or NMN-treated poor-grade oocytes, with excellent-grade oocytes serving as controls. Matured oocytes were subjected to in vitro fertilization (IVF), followed by embryo culture. **(W)** Representative images of embryos derived from excellent-grade oocytes and NMN-treated poor-grade oocytes following IVF and culture. **(X)** Quantification of the proportion of embryos reaching the eight-cell stage in the indicated groups. Data are presented as mean ± SD or SEM. The number of oocytes or embryos analyzed is indicated in the respective panels, where available. Data are presented as mean ± SEM. Statistical significance is indicated by asterisks (*P < 0.05, **P < 0.01, ***P < 0.001, ****P < 0.0001, n.s. = non-significant. Statistical significance was assessed using Mann-Whitney test **(K)** and unpaired t-test. Scale bars: 100 μm except 50 μm (L) and 10 μm **(O).**

We next examined whether these changes were accompanied by increased oxidative stress and alterations in mitochondrial status. ROS-associated fluorescence was higher in poor-grade oocytes than in excellent-grade oocytes (Intensity: 1042 ± 397 (s.d.) vs 540 ± 119 (s.d.); poor vs excellent; Fig. 5D,E), indicating an increased oxidative burden. MitoTracker Green fluorescence also differed between the two groups (Intensity: 860 ± 776 (s.d.) 1008 ± 921(s.d.); poor vs excellent; Fig. 5J,K), suggesting a difference in mitochondrial content or distribution. As this probe does not directly measure mitochondrial function, we additionally assessed mitochondrial membrane potential using JC-1. The red-to-green fluorescence ratio was markedly lower in poor-grade oocytes than in excellent-grade oocytes (Intensity: 0.64± 0.16 vs 1.28 ± 0.51; poor vs excellent; Fig. 5L,M), consistent with mitochondrial depolarization. Together, these observations reveal a coordinated phenotype in poor-grade oocytes involving altered ER-associated fluorescence, increased cytosolic and mitochondrial calcium signals, elevated ROS-associated fluorescence and reduced mitochondrial membrane potential.

Because proper meiotic spindle organization is essential for chromosome segregation and successful oocyte maturation, we next examined spindle morphology in excellent- and poor-grade oocytes. Oocytes were stained with an anti-α-tubulin antibody to visualize the meiotic spindle and classified into four morphological categories (Types I–IV; Fig. 5N,O). Excellent-grade oocytes predominantly displayed Type I spindle morphology, whereas poor-grade oocytes showed a marked shift towards Type IV morphology. Quantification showed that Type I spindles accounted for 46% of excellent-grade oocytes and 8.5% of poor-grade oocytes, whereas Type IV spindles accounted for 19% and 68%, respectively (Fig. 5P). Type II and Type III spindles represented 18% and 17% of excellent-grade oocytes, compared with 8.5% and 15% of poor-grade oocytes. This distribution indicates that poor oocyte quality is associated with a shift in spindle morphology, consistent with compromised organization of the meiotic apparatus. In the context of the calcium and mitochondrial abnormalities described above, these findings suggest that poor-grade oocytes experience defects across several cellular systems required for normal meiotic progression.

Having identified these abnormalities, we next asked whether restoring NAD⁺ availability could alleviate the altered cellular state of poor-grade oocytes. Based on our earlier observation that NMN supplementation improved meiotic maturation, we supplemented poor-grade oocytes with NMN during in vitro maturation (IVM) and reassessed ROS, calcium signals and mitochondrial membrane potential, using excellent-grade oocytes as the reference group (Fig. 5Q).

Following NMN supplementation, ROS-associated fluorescence in poor-grade oocytes was comparable to that in excellent-grade oocytes (8353 ± 4142 (s.d.) vs 7613 ± 2964 (s.d.); poor +NMN vs excellent; Fig. 5R & Fig. S5A). Similarly, Fluo-3 AM fluorescence did not differ significantly between the groups (548 ± 130 (s.d.) vs 513 ± 118 (s.d.); poor +NMN vs excellent; Fig. 5S & Fig. S5B), suggesting that NMN supplementation alleviated the increased cytosolic calcium signal. Rhod-2 AM fluorescence was also comparable between NMN-treated poor-grade oocytes and excellent-grade controls (730 ± 473 (s.d.) vs 848 ± 443 (s.d.); poor +NMN vs excellent; Fig. 5T & Fig. S5C), indicating an improvement in the mitochondrial calcium-associated signal. These observations suggest that NMN supplementation was accompanied by a reduction in oxidative stress and an improvement in calcium homeostasis.

We next assessed mitochondrial membrane potential following NMN treatment. The JC-1 red-to-green fluorescence ratio was higher in NMN-treated poor-grade oocytes than in the excellent-grade reference group comparison shown in Fig. 5U & Fig. S5D (0.95 ± 0.289 (s.d.) vs 0.45 ± 0.162 (s.d.); poor +NMN vs excellent) consistent with an improvement in mitochondrial polarization. Taken together, the NMN-associated changes in ROS, calcium signals and JC-1 fluorescence indicate that NAD⁺ precursor supplementation can alleviate several of the cellular abnormalities observed in poor-grade oocytes. However, because spindle morphology was assessed separately in Fig. 5N–P, these data do not establish whether NMN also restores spindle organization.

Finally, we investigated whether NMN supplementation was associated with improved developmental potential after fertilization. Following IVM, oocytes were subjected to in vitro fertilization and the resulting embryos were cultured to assess early development (Fig. 5V). Embryos derived from excellent-grade oocytes and NMN-treated poor-grade oocytes showed comparable morphological features (Fig. 5W). The proportion of embryos reaching the ≥8-cell stage was 60 % for excellent-grade oocytes and 56% for NMN-treated poor-grade oocytes, with no significant difference between the groups (Fig. 5X). These results indicate that NMN-treated poor-grade oocytes were able to support early embryonic development to a level comparable to that observed with excellent-grade oocytes under the conditions tested.

### NAD⁺ restoration promotes epigenetic remodeling and preserves oocyte competence through SIRT6-associated regulation

Above results showed that NMN supplementation improved meiotic maturation and alleviated several cellular abnormalities in poor-grade oocytes. Because NAD⁺ is required for the activity of sirtuins and other enzymes involved in chromatin regulation, we next investigated whether NAD⁺ restoration was associated with changes in the epigenetic state of poor-grade oocytes and whether SIRT6 contributed to the improvement in meiotic competence.

Comparative proteomic analysis revealed differences in the abundance profiles of multiple proteins associated with DNA methylation, histone modification and chromatin regulation between excellent- and poor-grade oocytes (Fig. 6A). These proteins included DNMT1, DNMT3A and DNMT3B, which are involved in DNA methylation, and TET1, which participates in DNA demethylation. Differences were also observed among histone-modifying enzymes, including KAT2A, KAT2B, HDACs, KMTs and KDMs. SIRT6, an NAD⁺-dependent histone deacetylase, was also represented in the differential proteomic profile. Together, these findings indicate that poor-grade oocytes differ from excellent-grade oocytes in the abundance of several components of the chromatin-regulatory machinery, raising the possibility that altered NAD⁺ metabolism is associated with changes in epigenetic regulation. To determine whether these proteomic differences were accompanied by changes in DNA methylation-associated fluorescence, we assessed 5-methylcytosine (5mC) in excellent- and poor-grade oocytes before IVM (Fig. 6D,E). Poor-grade oocytes exhibited higher 5mC fluorescence than excellent-grade oocytes (1399 ± 325 (s.d.) vs 811 ± 374 (s.d.); poor vs excellent; Fig. 6E), indicating an increased global 5mC-associated signal. We then examined whether NMN supplementation was associated with a change in this phenotype. Following IVM in the presence of NMN, 5mC fluorescence in poor-grade oocytes was no longer significantly different from that in excellent-grade oocytes (1463 ± 347 (s.d.) vs 1327 ± 234 (s.d.); poor+NMN vs excellent; Fig. 6F). These findings suggest that NMN supplementation is associated with a shift in the 5mC fluorescence profile of poor-grade oocytes towards that observed in excellent-grade controls. However, fluorescence-based 5mC measurements do not establish changes at specific genomic loci or demonstrate that all aspects of DNA methylation have been restored.

**Figure 6.**
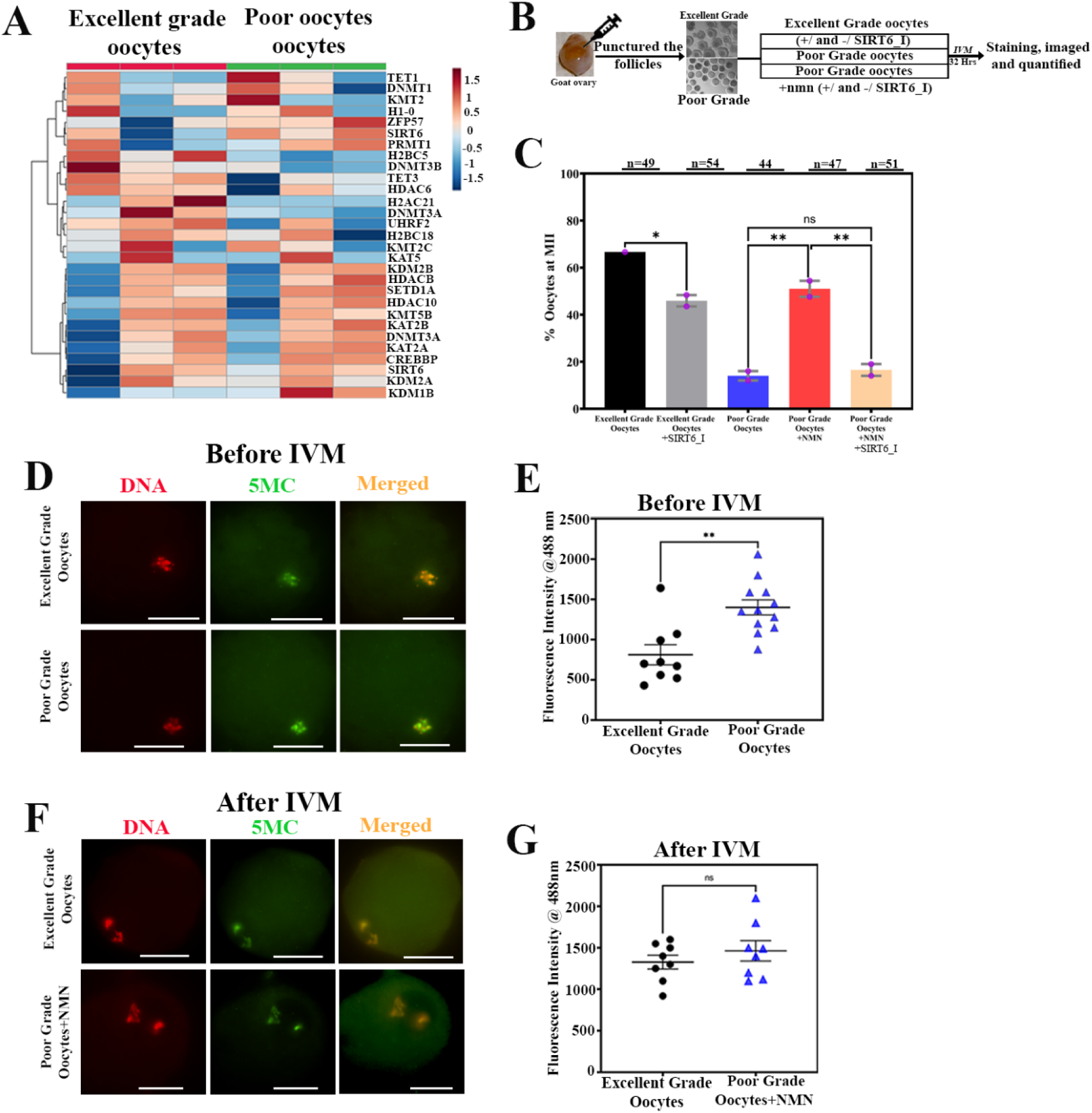
NAD⁺ restoration and SIRT6 inhibition reveal a functional association between NAD⁺ metabolism and epigenetic regulation in poor-grade goat oocytes. **(A)** Heatmap showing the relative abundance of selected proteins associated with epigenetic regulation in excellent-grade and poor-grade oocytes. Hierarchical clustering illustrates differences in the abundance patterns of chromatin-associated proteins and epigenetic regulators between the two groups. **(B)** Schematic of the experimental workflow for collecting goat oocytes, applying the indicated NMN and SIRT6 inhibitor treatments, and assessing meiotic maturation following in vitro maturation (IVM). **(C)** Quantification of the proportion of oocytes reaching metaphase II (MII) in the indicated experimental groups: excellent-grade controls, excellent-grade oocytes treated with a SIRT6 inhibitor, untreated poor-grade oocytes, NMN-treated poor-grade oocytes, and NMN-treated poor-grade oocytes exposed to a SIRT6 inhibitor. **(D)** Representative fluorescence images showing DNA staining (red), 5-methylcytosine (5mC; green), and merged images in excellent-grade oocytes and untreated poor-grade oocytes; **(E)** Quantification of 5mC-associated fluorescence intensity before IVM in excellent-grade and poor-grade oocytes **(F)** Representative fluorescence images showing DNA staining (red), 5-methylcytosine (5mC; green), and merged images in excellent-grade oocytes and NMN-treated poor-grade oocytes. **(G)** Quantification of 5mC-associated fluorescence intensity following IVM in excellent-grade oocytes and NMN-treated poor-grade oocytes (F). Data are presented as mean ± SD or SEM. The number of oocytes analyzed is indicated above the respective bars. Statistical significance is indicated by asterisks (*P < 0.05, **P < 0.01, ***P < 0.001, ****P < 0.0001, n.s. = non-significant. Statistical significance was assessed using ordinary one-way ANOVA **(C)** and unpaired t-test. Scale bars: 50 μm.

The differential representation of SIRT6 in the proteomic profile prompted us to investigate whether SIRT6 activity contributes to the beneficial effect of NMN on meiotic maturation. We compared maturation outcomes in excellent- and poor-grade oocytes following IVM with NMN supplementation and/or SIRT6 inhibition (Fig. 6B,C & Fig. S6A). As expected, excellent-grade oocytes showed a higher maturation rate than untreated poor-grade oocytes, with 66.65% and 14.0 ± 2.0% (s.e.m.) reaching metaphase II (MII), respectively (Fig. 6C & Fig. S6A). NMN supplementation increased the proportion of poor-grade oocytes reaching MII from 14.0 ± 2.0% (s.e.m.) to 51 ± 3.4% (s.e.m.). However, when SIRT6 was inhibited in the presence of NMN, the maturation rate decreased to 16.5 ± 2.5% (s.e.m.), substantially weakening the NMN-associated improvement. SIRT6 inhibition also reduced the MII rate in excellent-grade oocytes from 66.7% to 45.9 ± 2.42% (s.e.m.) (Fig. 6C & Fig. S6A). These findings indicate that SIRT6 activity contributes to normal meiotic maturation and is required, at least in part, for the full maturation benefit associated with NMN supplementation under the conditions tested.

### NMN supplementation improves meiotic competence in poor-grade oocytes across large mammalian species

Given NMN supplementation improved meiotic maturation in poor-grade goat oocytes, we next investigated whether this beneficial effect extended to other large mammalian species. Bovine and porcine oocytes were selected to determine whether the response to NMN was reproducible across distinct livestock models (Fig. 7).

**Figure 7.**
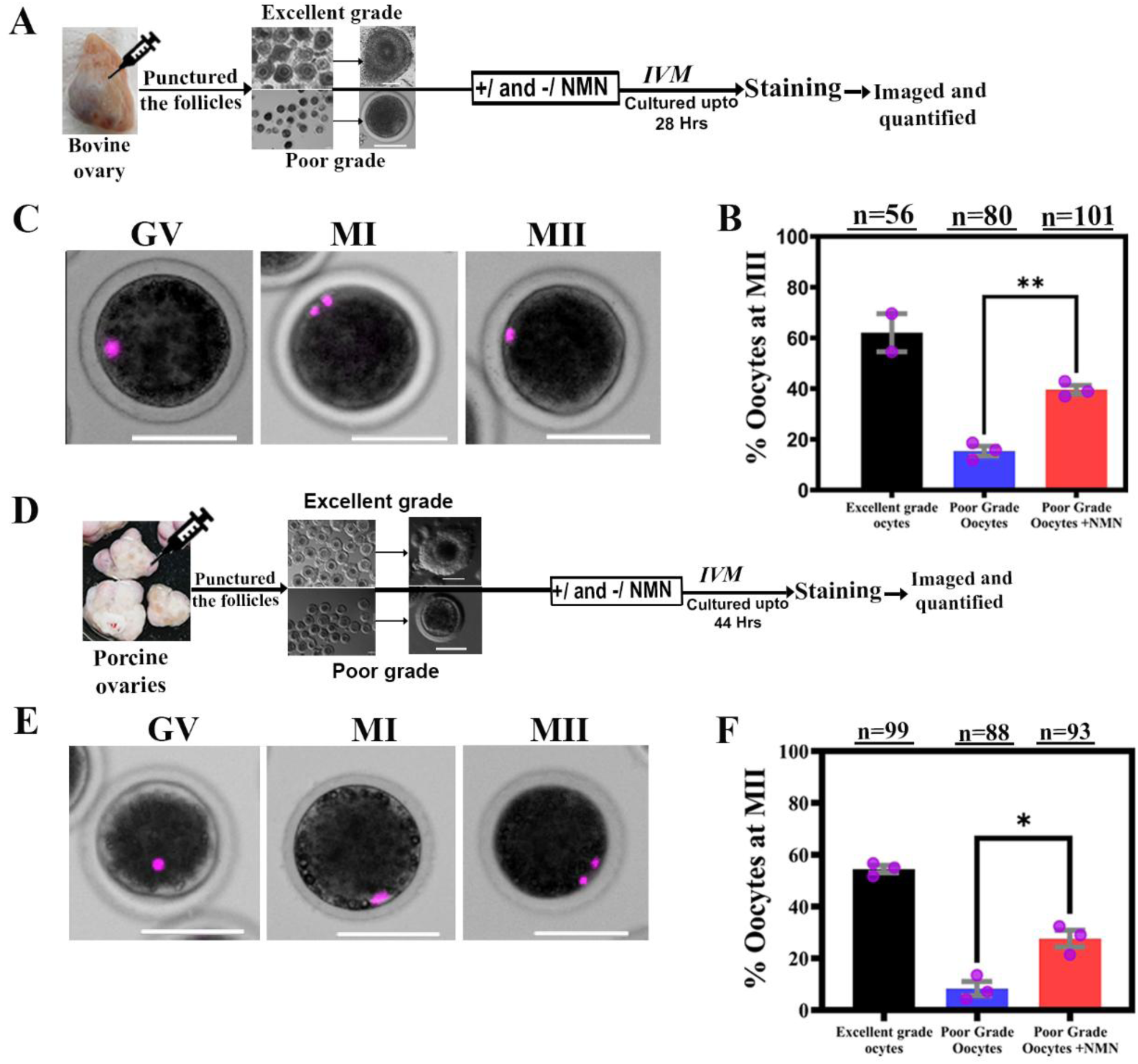
NMN supplementation improves meiotic maturation in poor-grade bovine and porcine oocytes. **(A)** Schematic of the experimental workflow for bovine oocytes, including follicle aspiration, oocyte collection, NMN supplementation during in vitro maturation (IVM), and subsequent staining, imaging, and quantification. **(B)** Representative images of bovine oocytes at the germinal vesicle (GV), metaphase I (MI), and metaphase II (MII) stages following DNA staining with DAPI (magenta). **(C)** Quantification of the proportion of bovine oocytes reaching metaphase II (MII) following IVM in excellent-grade oocytes, untreated poor-grade oocytes, and NMN-treated poor-grade oocytes. The number of oocytes analyzed in each group is indicated above the bars. **(D)** Schematic of the experimental workflow for porcine oocytes, including follicle aspiration, oocyte collection, NMN supplementation during IVM, and subsequent staining, imaging, and quantification. **(E)** Representative images of porcine oocytes at the germinal vesicle (GV), metaphase I (MI), and metaphase II (MII) stages following DNA staining with DAPI (magenta). **(F)** Quantification of the proportion of porcine oocytes reaching MII following IVM in excellent-grade oocytes, untreated poor-grade oocytes, and NMN-treated poor-grade oocytes. The number of oocytes analyzed in each group is indicated above the bars. Data are presented as mean ± SEM. Statistical significance is indicated by asterisks (*P < 0.05, **P < 0.01, ***P < 0.001, ****P < 0.0001. Statistical significance was assessed using ordinary one-way ANOVA. Scale bars: 100 μm

We first examined bovine oocytes collected from punctured follicles and classified according to morphological quality before in vitro maturation (IVM; Fig. 7A & Fig. S7A). As observed in goat oocytes, poor-grade bovine oocytes exhibited substantially lower meiotic competence than excellent-grade oocytes, with the proportion reaching metaphase II (MII) decreasing from 62.05 ± 7.51% (s.e.m.) to 15.35 ± 1.96% (s.e.m.) (Fig. 7B & Fig. S7B). NMN supplementation increased the MII rate in poor-grade bovine oocytes from 15.35 ± 1.96% (s.e.m.) to 39.58 ± 1.73% (s.e.m.), demonstrating a significant improvement in meiotic maturation. Although the response did not fully restore maturation to the level observed in excellent-grade oocytes, the improvement supports the potential of NAD⁺ precursor supplementation to enhance the competence of poor-grade bovine oocytes.

We next examined porcine oocytes to determine whether a similar response occurred in another large mammalian species (Fig. 7C & Fig. S7C). Poor-grade porcine oocytes also showed markedly reduced meiotic maturation compared with excellent-grade oocytes, with MII rates of 8.3 ± 2.71% (s.e.m.) and 54.4 ± 1.45% (s.e.m.), respectively (Fig. 7D & Fig. S7D). Supplementation with NMN increased the proportion of poor-grade porcine oocytes reaching MII from 8.3 ± 2.71% (s.e.m.) to 27.6 ± 3.23% (s.e.m.), indicating a significant improvement in maturation. Although the extent of improvement differed between the bovine and porcine models, the response was consistent in direction across both species. Together with our findings in goat oocytes, these results demonstrate that NMN supplementation improves the meiotic maturation of poor-grade oocytes in three large mammalian species. The reproducibility of this response across goat, cattle and pigs suggests that the beneficial effect of NAD⁺ precursor supplementation is not restricted to a single species. These findings extend the translational relevance of our study and support further investigation of NAD⁺ metabolism as a potential target for improving the utilization of poor-grade oocytes in livestock reproductive technologies. However, these experiments establish a cross-species effect of NMN on meiotic maturation; they do not independently demonstrate that the QPRT–NAD⁺ pathway or the downstream mitochondrial and epigenetic mechanisms identified in goat oocytes operate identically in bovine and porcine oocytes.

## Discussion

Oocyte developmental competence is usually judged from morphology, particularly cumulus investment and cytoplasmic appearance, yet these features provide only an indirect indication of the physiological state of the oocyte^32,33^. Our findings suggest that poor oocyte quality reflects a deeper metabolic and stress-associated state that is already established before meiotic maturation. Poor-grade goat oocytes showed substantially reduced meiotic competence, and this defect persisted despite additional cumulus-cell support. This distinction is important because cumulus cells provide metabolites, substrates and regulatory signals required for oocyte maturation, while the oocyte itself has a distinct metabolic program that depends heavily on mitochondrial metabolism^12^. Thus, although an intact cumulus–oocyte interaction is important for competence, our experiments and previous reports indicate that the phenotype of poor-grade oocytes cannot be explained simply by insufficient somatic-cell support. Instead, the defect appears to reside, at least in substantial part, within the oocyte itself.

Metabolomic profiling provided an initial clue to the nature of this intrinsic defect. Excellent- and poor-grade GV oocytes occupied clearly distinct metabolic states, with NAD⁺ metabolism emerging as one of the most prominent differences. Poor-grade oocytes had reduced NAD⁺ together with accumulation of quinolinic acid and depletion of downstream NAD⁺-biosynthetic intermediates. This pattern pointed specifically to the conversion of quinolinic acid to nicotinic acid mononucleotide (NAMN), catalysed by quinolinate phosphoribosyltransferase (QPRT), rather than to a generalized limitation in precursor availability. Consistent with this interpretation, tryptophan uptake was not impaired and additional tryptophan failed to improve maturation. In contrast, bypassing QPRT with NAMN or restoring QPRT itself improved meiotic maturation. Together, these experiments identify QPRT-dependent de novo NAD⁺ synthesis as a functional metabolic bottleneck in poor-quality oocytes.

The importance of this pathway is supported by genetic evidence from mice. Deletion of enzymes involved in de novo NAD⁺ synthesis, including Qprt, lowers ovarian NAD⁺ and is associated with loss of ovarian reserve, impaired oocyte quality, spindle abnormalities, increased oxidative stress and reduced fertilization and early embryonic development^25^. Qprt-deficient ovaries also show extensive transcriptional changes involving mitochondrial metabolic pathways, accompanied by abnormalities in mitochondrial distribution and membrane potential. Importantly, several of these defects can be ameliorated by NAD⁺ precursor supplementation^25^. These observations provide an independent genetic framework for our findings and indicate that disruption of QPRT-dependent NAD⁺ metabolism is sufficient to compromise ovarian and oocyte function^25^.

Our data further provide a potential explanation for how this metabolic defect develops in poor-quality oocytes. Proteomic analysis identified pronounced changes in pathways related to ER function, protein homeostasis and the unfolded protein response, accompanied by activation of IRE1α signalling and increased XBP1 splicing. Manipulating ER stress altered meiotic competence in both directions: induction of ER stress compromised maturation of otherwise excellent-grade oocytes, whereas attenuation of ER stress improved maturation of poor-grade oocytes. Inhibition of IRE1α similarly improved maturation. ER stress therefore appears not simply to accompany poor oocyte quality, but to contribute functionally to the loss of competence. The connection between ER stress and QPRT is particularly intriguing. Previous work in kidney injury demonstrated that impairment of de novo NAD⁺ synthesis can occur as an early component of the ER-stress response. In that system, ER stress suppressed QPRT expression, resulting in defective quinolinate utilization and impaired NAD⁺ biosynthesis; the response involved the ER-stress transcription factor DDIT3/CHOP^31^. Thus, regulation of QPRT may represent a broader point of intersection between cellular stress and NAD⁺ metabolism rather than an oocyte-specific phenomenon. Our findings extend this relationship to oocytes but suggest that the underlying regulation may differ. QPRT RNA declined more rapidly in poor-grade oocytes during RNA-chase experiments, and recombinant IRE1α was capable of cleaving QPRT RNA in vitro. Together with increased IRE1α activity in poor-grade oocytes, these observations raise the possibility that IRE1α-dependent RNA processing contributes to QPRT loss.

IRE1α provides a plausible molecular link between ER stress and the disruption of QPRT-dependent NAD⁺ biosynthesis in poor-grade oocytes. In addition to its canonical role in XBP1 mRNA splicing, IRE1α can regulate cellular RNA homeostasis through regulated IRE1-dependent decay (RIDD), targeting selected transcripts for cleavage. In our study, increased IRE1α activity in poor-grade oocytes coincided with reduced QPRT RNA stability, while recombinant IRE1α directly cleaved QPRT RNA in vitro. These complementary observations support a model in which ER stress-associated IRE1α activation contributes to QPRT transcript destabilization, potentially through a RIDD-like mechanism. This provides a mechanistic explanation for the reduced QPRT expression and consequent impairment of de novo NAD⁺ biosynthesis observed in poor-grade oocytes. Defining the endogenous QPRT cleavage site and establishing its dependence on IRE1α activity will further delineate this regulatory mechanism in oocytes. Nevertheless, our findings identify QPRT RNA as a potential target of IRE1α-mediated regulation and establish a functional connection between ER stress signalling and NAD⁺ metabolic insufficiency.

The consequences of disturbed NAD⁺ homeostasis extended well beyond NAD⁺ metabolism itself. Poor-grade oocytes showed abnormal ER organization, altered cytosolic and mitochondrial Ca²⁺, increased oxidative stress, loss of mitochondrial membrane potential and disturbed mitochondrial distribution. These processes are mechanistically interconnected. The ER and mitochondria coordinate cellular Ca²⁺ handling, while excessive mitochondrial Ca²⁺ loading can increase electron leakage and ROS generation and compromise mitochondrial polarization. NAD⁺ availability, in turn, influences redox balance, mitochondrial metabolism and NAD⁺-dependent signalling. The simultaneous disruption of these processes therefore suggests that NAD⁺ depletion occurs within a broader failure of metabolic and organelle homeostasis rather than acting as an isolated metabolic lesion.

The rescue experiments support this interpretation. NMN supplementation did not merely increase the proportion of oocytes reaching MII; it reduced oxidative stress, improved mitochondrial membrane potential and partially normalized Ca²⁺ homeostasis. Importantly, the benefit extended beyond nuclear maturation to early embryonic development. This distinction is critical because polar-body extrusion alone does not necessarily indicate acquisition of developmental competence. Restoration of post-fertilization development suggests that replenishing NAD⁺ improves the underlying physiological quality of the oocyte rather than simply permitting meiotic progression.

NAD⁺ availability may also connect metabolic state with chromatin regulation^34^. Proteomic analysis identified changes in multiple chromatin-associated and epigenetic regulators, including NAD⁺-responsive proteins, while poor-grade oocytes showed altered global DNA methylation during maturation. Inhibition of SIRT6 attenuated the beneficial effect of NMN, suggesting that at least part of the response to NAD⁺ restoration requires NAD⁺-dependent signalling rather than being explained solely by increased metabolic substrate availability. This provides a potential route through which metabolic state could influence the chromatin transitions required for acquisition of developmental competence. We do not infer from these experiments that SIRT6 alone accounts for the chromatin phenotype; rather, they place NAD⁺-dependent chromatin regulation downstream of the metabolic defect identified in poor-quality oocytes.

The ability of NMN to improve maturation in poor-quality bovine and porcine oocytes further suggests that this relationship is not restricted to the goat. This cross-species conservation is particularly relevant because morphological heterogeneity among oocytes is a common feature of assisted reproduction and livestock embryo production. Previous studies have already shown that increasing NAD⁺ availability can improve oocyte quality, mitochondrial function and fertility in experimental models. For example, NMN supplementation restores several aspects of oocyte function in reproductively aged mice, while manipulation of NMNAT1/3 or NAMPT influences meiotic progression and spindle organization^20,35^. Our study adds a mechanistic dimension to these observations by identifying a QPRT-associated defect in endogenous NAD⁺ production and linking it to cellular stress in intrinsically poor-quality oocytes.

To investigate the translational relevance of the relationship between ER proteostasis and NAD⁺ metabolism identified in goat oocytes, we further analysed published proteomic data from human germinal vesicle (GV)-stage oocytes obtained from young and advanced-maternal-age (AMA) women (Fig. 8A–C)^36^. Principal component analysis revealed separation between the two age groups, suggesting that maternal age is associated with a distinct oocyte proteomic state (Fig. 8B). Examination of ER proteostasis-associated proteins revealed increased abundance of HSPA5, SIL1 and PDIA3 in AMA oocytes, with nominal P values of 0.031, 0.007 and 0.02, respectively (Fig. 8C). These proteins participate in ER protein folding and quality control, suggesting that the ER proteostasis machinery is remodelled in human oocytes with advancing maternal age. However, increased abundance of these proteins does not, by itself, establish activation of the unfolded protein response or demonstrate ER dysfunction; it may also reflect a compensatory response to increased proteostatic demand.

**Figure 8.**
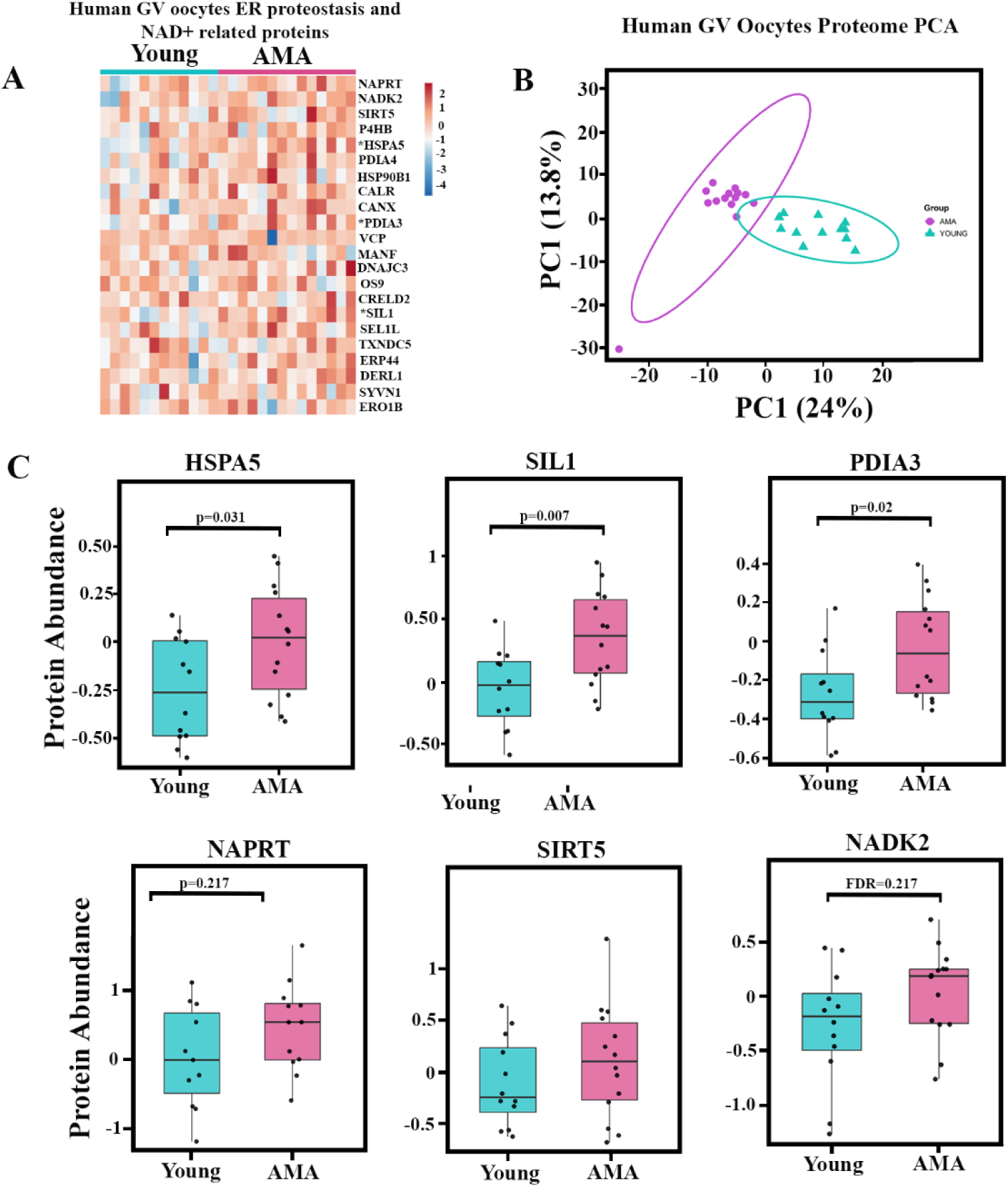
Comparative proteomic profiling of human germinal vesicle (GV) oocytes from young and advanced maternal age (AMA) groups. **(A)** Heatmap showing the relative expression levels of selected proteins in young and advanced maternal age (AMA) human germinal vesicle (GV) oocytes. Colours represent relative protein expression, with different colors indicating higher and lower expression levels following row-wise Z-score normalization. **(B)** Principal component analysis (PCA) showing the distribution of proteomics profiles between Young and AMA human germinal vesicle (GV) oocytes. PC1 and PC2 explain 24% and 13.3% of the total variance, respectively. **(C)** Boxplots comparing the expression levels of *HSPA5*, *SIL1*, *PDIA3*, *NAPRT*, *SIRT5*, and *NADK2* between Young and AMA human GV oocytes. *HSPA5* (heat shock protein family A member 5), *SIL1* (SIL1 nucleotide exchange factor), and *PDIA3* (protein disulfide isomerase family A member 3) are associated with ER stress and protein folding, whereas *NAPRT* (nicotinate phosphoribosyltransferase), *SIRT5* (sirtuin 5), and *NADK2* (NAD kinase 2) are associated with NAD metabolism. Each point represents an individual sample. Statistical significance was assessed using an unpaired t-test.

We also examined NAD-related proteins in the same human dataset. NAPRT, NADK2 and SIRT5 showed trends towards altered abundance in AMA oocytes (Fig. 8C). Although these changes did not remain statistically significant after correction for multiple testing, their pattern raises the possibility that age-associated changes in ER proteostasis occur alongside remodelling of NAD-related metabolic and mitochondrial regulatory pathways. Importantly, protein abundance alone cannot establish intracellular NAD⁺ levels, NAD⁺/NADH balance or the activity of individual biosynthetic pathways. These findings therefore indicate coordinated proteomic changes rather than demonstrating NAD⁺ deficiency in human AMA oocytes. The human findings complement, but are biologically distinct from, our comparison of excellent-grade and poor-grade goat oocytes (Fig. 8D). Whereas the goat model identifies molecular differences associated with naturally occurring variation in oocyte competence, the human dataset examines changes associated with advanced maternal age. In goat oocytes, our functional experiments link ER stress and IRE1α signalling to reduced QPRT expression, impaired de novo NAD⁺ biosynthesis and compromised meiotic competence. The human proteomic data provide an independent translational context in which ER proteostasis and NAD-related proteins are altered with maternal age, but do not establish that the same QPRT-dependent mechanism operates in human oocytes. Together, these complementary datasets support further investigation of the relationship between ER homeostasis and NAD⁺ metabolism as a potential determinant of mammalian oocyte competence and reproductive aging.

Taken together, our findings support a model in which poor oocyte competence arises from an interconnected failure of stress adaptation and metabolic homeostasis. In poor-quality goat oocytes, ER stress and IRE1α activation are associated with reduced QPRT function and impaired de novo NAD⁺ biosynthesis. The resulting disturbance in NAD⁺ homeostasis is accompanied by altered Ca²⁺ handling, mitochondrial dysfunction, oxidative stress and changes in chromatin regulation, collectively limiting meiotic and developmental competence. Bypassing the metabolic bottleneck or replenishing NAD⁺ reverses several of these defects and restores competence, with NMN-mediated rescue extending across goat, bovine and porcine oocytes. Mining of human oocyte proteome independently identifies coordinated remodeling of ER-stress and NAD⁺ metabolic programs with advanced maternal age, providing a translational link to human reproductive biology. Rather than viewing morphologically poor oocytes simply as irreversibly damaged cells, these results suggest that at least a subset occupies a metabolically compromised but recoverable state. This concept has practical implications for both reproductive biology and assisted reproduction: intrinsic oocyte quality may be determined not only by the extent of accumulated cellular damage, but also by whether metabolic and stress-response networks retain sufficient plasticity to be restored. Defining the mechanisms that govern this plasticity and determining whether NAD⁺ homeostasis can be safely manipulated in human oocytes will be important before these findings can be translated into strategies for improving oocyte developmental competence.

## Abbreviation

FBS: fetal bovine serum
BSA: Bovine serum albumin
HEPES: N-[2-Hydroxyethyl]Piperazine-N[2-Ethane Sulphonic Acid]
FSH: Follicle-stimulating hormone
LH: Luteinizing hormone
hEGF: Human epidermal growth factor
hMG: Human menopausal gonadotropin
DMSO: Dimethyl sulfoxide
NMN: Nicotinamide mononucleotide
QA: Quinolinic acid
NAMN: Nicotinic acid mononucleotide
TUDCA: Tauroursodeoxycholic acid
SDC: Sodium deoxycholate
AMBIC: Ammonium bicarbonate
ROS: Reactive oxygen species
H₂DCFDA: 2′,7′-Dichlorodihydrofluorescein diacetate
PFA: Paraformaldehyde
5mC: 5-Methylcytosine
HCl: Hydrochloric Acid
DTT: Dithiothreitol

## Materials and Methods

### Animal Ethics

Goat ovaries, bovine and porcine ovaries were obtained from abattoir after slaughtering animals for commercial reasons. No animals were explicitly sacrificed for this particular study. The research was performed in conformity with the appropriate institutional standards on the use of animal tissues from slaughterhouses.

All procedures were approved by the Committee for Control and Supervision of Experiments on Animals (CCSEA) (V-11011(13)/11/ 2020-CPCSEA-DADF) and the Institutional Animal Ethics Committee (IAEC) (IAEC/2019/NIAB/10/HBDPR and IAEC/NIAB/2025/02/HBDPR) and the institutional Bio-Safety Committee (IBSC/2018/NIAB/HBDPR/001 and IBSC/April/2024/NIAB/PR009).

### Goat oocyte collection and morphological classification

Osmanabadi goat ovaries were obtained from a local abattoir and transported to the laboratory in pre-warmed 1× phosphate-buffered saline (PBS) containing penicillin–streptomycin (Sigma Aldrich, P4333) at 37°C using an embryo transport thermobox. Ovaries were processed within 2 h of collection. After removal of excess tissue, the ovaries were washed 7–8 times with pre-warmed PBS containing penicillin–streptomycin. Ovarian follicles were punctured using a 22-gauge needle into TCM-199 (Sigma Aldrich, M4530) supplemented with 5% fetal bovine serum (FBS) (ThermoFisher, 10270106), 50 µg/mL penicillin–streptomycin, and 18 mM HEPES, pH 7.25. Cumulus–oocyte complexes (COCs) were recovered under a stereomicroscope and classified according to their morphology. Excellent-grade oocytes were characterized by a compact cumulus consisting of more than three layers and homogeneous cytoplasm. Poor-grade oocytes were characterized by sparse or absent cumulus cells and heterogeneous, dark or granular cytoplasm. Oocytes selected for experiments were washed 5–6 times in the collection medium before further processing. Before being used, all media used in the experiments were filtred using a 0.22 µm syringe filter (TPP, 99722) equilibrated at 37°C under 5% CO₂ in humidified air for at least three hours.

### In vitro maturation of goat, bovine and porcine oocytes

Goat oocytes were matured in 150-µL microdrops of maturation medium under mineral oil (Sigma Aldarich, M8410). The maturation medium consisted of TCM-199 supplemented with 8 IU/mL follicle-stimulating hormone (FSH, Sigma Aldrich, F4021), 4 IU/mL luteinizing hormone (LH, Sigma Aldrich, L6420), 1 µg/mL estradiol-17β, 10% goat follicular fluid, 10% FBS, 0.81 mM sodium pyruvate (Sigma Aldrich, P4562), 10 ng/mL human epidermal growth factor (hEGF), and 50 µg/mL gentamicin. Oocytes were cultured for 32 h at 38.5°C in a humidified incubator containing 5% CO₂.

Bovine and porcine oocytes were matured using the same basic maturation system with species-specific modifications. For bovine oocytes, 10% bovine follicular fluid was used instead of goat follicular fluid, and 10 IU/mL human menopausal gonadotropin (hMG, Sanzyme) was used instead of FSH and LH; cultured for 28 h. For porcine oocytes, 10% porcine follicular fluid was used in place of goat follicular fluid and cultured for 44 h while the remaining components of the maturation medium were maintained as described above.

Unless otherwise specified, all chemical treatments were added directly to the maturation medium and compared with untreated or vehicle-treated controls under otherwise identical maturation conditions.

### Assessment of cumulus-cell contribution to oocyte maturation

To determine whether the maturation defect of poor-grade oocytes could be influenced by cumulus-cell support, excellent- and poor-grade oocytes were cultured either with their original cumulus cells or following removal of cumulus cells. For cumulus removal, COCs were incubated with 0.2 mg/mL hyaluronidase (Sigma Aldrich, H3506) for 5 min at 37°C and gently pipetted until the cumulus cells were removed. For the cumulus-cell supplementation experiment, poor-grade oocytes were cultured in the presence of additional cumulus cells bedding under standard IVM conditions. Following 32 h of maturation, nuclear maturation was assessed as described below.

### Chemical treatments during in vitro maturation

A series of supplementation and pharmacological intervention experiments was performed during IVM to examine the effects of NAD⁺ metabolism and ER stress on meiotic maturation.

### Quercetin treatment

Quercetin (SRL, 71923) was used to examine whether reducing NAD⁺ consumption could improve the maturation of poor-grade oocytes. A 1 mM stock solution was prepared in DMSO (SRL, 66444) and stored at −20°C. The stock was diluted in IVM medium to a final concentration of 50 µM. Poor-grade oocytes were matured in the presence of quercetin or the corresponding concentration of DMSO for 32 h.

### NMN supplementation

Nicotinamide mononucleotide (NMN) (SRL, 95130) was used to increase NAD⁺ precursor availability. A 10 mM stock solution was prepared in TCM-199 and stored at −20°C. For goat oocytes, NMN was added to the IVM medium at a final concentration of 400 µM. Porcine poor-grade oocytes were treated with 550 µM NMN. Oocytes were matured under the standard conditions described above.

### Tryptophan supplementation

To examine whether substrate availability influenced de novo NAD⁺ biosynthesis, poor-grade oocytes were supplemented with exogenous 750 µm L-tryptophan (Sigma Aldrich, T0254) during IVM. The concentration used for the experiment should be specified according to the experimental record.

### Quinolinate and NAMN supplementation

Quinolinic acid (QA) (Cayman Chemicals, 114941-1 gm) 600 µm and nicotinic acid mononucleotide (NAMN) (Cayman Chemicals, 32883-1 mg) 400 µm were added to the IVM medium to examine the effect of metabolites positioned upstream and downstream of the QPRT-dependent step of NAD⁺ biosynthesis.

### TUDCA treatment

Tauroursodeoxycholic acid (TUDCA) (Combi-Blocks, QB-9028) was used to attenuate ER stress. A 200 mM stock solution was prepared in DMSO and stored at −20°C. For maturation experiments, TUDCA was added to the IVM medium at a final concentration of 200 µM. Poor-grade oocytes were cultured under standard IVM conditions for 32 h. For experiments examining QPRT expression, poor-grade oocytes were treated with 200 µM TUDCA for 14 h.

### STF-083010 treatment

STF-083010 (MedChemExpress, HY-15845) was used to inhibit IRE1α activity. A stock solution was prepared in fresh anhydrous DMSO and stored at −80°C. The final concentration used during IVM was 30 µM. Poor-grade oocytes were cultured under standard IVM conditions in the presence or absence of STF-083010.

### Tunicamycin treatment

Tunicamycin (Calbiochem, 654380) was used to induce ER stress in excellent-grade oocytes. A 5 mg/mL stock solution was prepared in DMSO and stored at −20°C. For IVM experiments, 0.1 µL of the stock solution was added to 100 µL of maturation medium to obtain a final concentration of 5 µg/mL. Excellent-grade oocytes were cultured under standard IVM conditions for 32 h.

### SIRT6-IN-1 Treatment

To assess the influence of the SIRT6-IN-1 (Sigma, Aldrich, SML3805) inhibitor in the oocytes, a 1 mM stock solution of SIRT6-IN-1 was prepared in DMSO and stored at −20 °C. Prior to use, it was diluted to a 100 µM concentration with the corresponding IVM media.

Excellent grade oocytes with and without SIRT6-IN-1, poor grade with NMN and SIRT6-IN-1, poor grade oocytes with and without NMN were kept for maturation under the standard conditions described above.

### Evaluation of nuclear maturation

Following IVM, cumulus cells were removed by incubation with 0.2 mg/mL hyaluronidase for 5 min at 37°C followed by gentle pipetting. Denuded oocytes were stained with Hoechst 33342 in TCM-199 for 10 min at 37°C and washed three times with PBS. The nuclear stage was assessed by fluorescence microscopy, and oocytes were classified according to meiotic stage. The proportion of oocytes reaching metaphase II (MII) was used as the primary endpoint of meiotic maturation.

### Fluorescent tryptophan uptake assay

Intracellular tryptophan availability was assessed using fluorescently labelled Fmoc-Trp-BODIPY (Sigma Aldrich, 900606). Excellent- and poor-grade oocytes were denuded and incubated with 20 µM Fmoc-Trp-BODIPY in IVM media for up to 40 minutes at 37°C.

Following incubation, oocytes were washed and imaged by fluorescence microscopy at 488 nm, and intracellular fluorescence intensity was quantified.

### NAD+ measurement

We collected 200 oocytes for each group in order to determine the NAD+ in excellent and poor grade oocytes. Following the manufacturer’s instructions, we used the NAD/quantitation kit (MAK037; Sigma-Aldrich) to assess the intracellular levels of NAD+.

### Untargeted metabolomic analysis

#### Extraction of follicular fluid metabolites using the MeOH/H₂O method

For untargeted metabolomic profiling, 250 denuded excellent-grade and 250 denuded poor-grade oocytes were collected for each biological replicate, with three independent biological replicates analyzed. Samples were snap-frozen and stored at −80°C until extraction. Oocytes were extracted using 200 µL of methanol/water (4:1, v/v). Samples were sonicated for 2 min using 5-s on/5-s off cycles at 40% amplitude on ice and incubated on ice for 20 min. Samples were centrifuged at 15,000 × g for 10 min at 4°C, and the resulting supernatants were collected and dried using a vacuum concentrator. Dried extracts were stored at −80°C until analysis. For LC–MS analysis, dried extracts were reconstituted in 100 µL of acetonitrile/water (3:1, v/v) and centrifuged at 14,000 rpm for 15 min at 4°C before transfer to LC vials.

### Instrumentation

Metabolomic analysis was performed using a Waters Xevo G2-XS QToF mass spectrometer coupled to a BEH C18 column (Xevo G2-XS QToF, Waters Corp., Milford, MA, USA). Water containing 0.1% formic acid was used as solvent A and methanol containing 0.1% formic acid as solvent B. The flow rate was 0.3 mL/min, and the total chromatographic run time was 35 min. Separate chromatographic gradients were used for the lipid and methanol fractions. The gradient elution program was set as follows: 85% A at 0–2 min, decreasing to 20% A at 6 min, 10% A at 10 min, 5% A at 15 min, and 1% A at 25 min. The mobile phase was then returned to the initial composition of 85% A at 31 min and maintained until 35 min for column re-equilibration. For the methanol fraction, the composition was 99% at 0–2 min, decreasing to 80% at 6 min, 40% at 10 min, 10% at 15 min, and 1% at 25 min. The initial composition of 99% was restored at 31 min and maintained until 35 min. Mass spectra were acquired in MSE mode using alternating low- and high-energy conditions over an m/z range of 50–2000 in both positive- and negative-ion modes. Collision energies of 15–40 V were used for high-energy acquisition. The capillary voltage was 2.0 kV, cone voltage 40 V, desolvation temperature 350°C, the desolvation gas flow 800 L/h, the cone gas flow 50 L/h and source temperature 120°C. Leucine enkephalin (Leu-Enk, m/z 554.262, [M − H]-) was used as the lock-mass reference.

### Metabolite analysis

Metabolomic data from excellent and poor grade oocyte samples was collected using positive and negative electrospray ionization mode. ProteoWizard software was used to process the raw data for format conversion, followed by MZmine 2.53 for mass spectrometric data analysis. MZmine was used for centroid-based mass detection, peak detection, alignment, and identification, outputting data as m/z ratios. Metabolite quantification using peak height intensity. Statistical significance (p-values) was computed using a t-test in Excel after log2-transforming intensity. Metabolites were identified using a 5-ppm mass tolerance in HMDB and LIPIDMAPS databases. The peak intensities were entered into MetaboAnalyst 6.0, filtered with a 40% standard deviation threshold and log2-transformed using Pareto scaling. We used Principal Component Analysis (PCA) to analyse the differences between oocytes groups, pathway enrichment was performed. The top 50 differently regulated metabolites between excellent and poor grade oocytes were identified using VIP plots (VIP score >1.3 for heatmap), fold-change graphs, and heatmaps. Pathway analysis was performed on control and cystic samples using MetaboAnalyst 6.0.

### Targeted metabolite analysis by HPLC

For targeted analysis of NAD⁺ metabolism, oocytes were collected separately according to morphological grade, denuded of cumulus cells, and processed for metabolite extraction. The extraction of metabolites was done by using MeOH/H₂O as described above. NAD⁺ and metabolites associated with the de novo NAD⁺ biosynthetic pathway, including quinolinic acid (QA), nicotinic acid mononucleotide (NAMN), and nicotinamide mononucleotide (NMN), were quantified by HPLC. NMN (nicotinamide mononucleotide), NAMN (nicotinic acid mononucleotide), and quinolinic acid were analyzed using a high-performance liquid chromatography (HPLC) system equipped with a UV detector (SPD-20A) operated by LC-solution software. Separation was performed using a C18 reversed-phase column (250 × 4.6 mm, 5 μm particle size) maintained at 30–35°C. The mobile phase consisted of 0.1% formic acid in water (Mobile Phase A) and 0.1% formic acid in methanol (Mobile Phase B). Gradient elution was performed at a flow rate of 1.0 mL/min using the following program: 98% A and 2% B from 0 to 5 min, 80% A and 20% B at 15 min, 50% A and 50% B at 20 min, followed by re-equilibration to 98% A and 2% B at 25 min. The initial mobile-phase composition was maintained until 30 min. Detection was performed at 260 nm, with a total run time of 30 min. NMN, NAMN, and quinolinic acid were quantified by comparing their chromatographic peak areas with those of the corresponding reference standards. A sample injection volume of 10 μL was used for each run.

### Proteomic analysis

#### Protein extraction and digestion

To compare the proteomes of excellent- and poor-grade oocytes, 200 completely denuded oocytes from each group were collected for each biological replicate. Three independent biological replicates were analyzed. Oocytes were lysed in 2% sodium deoxycholate (SDC, Sigma Aldrich, D6750) prepared in 50 mM ammonium bicarbonate (NH₄HCO₃, Sigma Aldrich, 5.33005), pH 7.2. Samples were sonicated for 2 min using 5-s on/5-s off cycles at 40% amplitude on ice and immediately heated at 95°C for 10 min. Protein concentration was determined using the Bradford Protein assay Kit (Takara, T9310A) according to the manufacturer’s instructions. Equal amounts of protein were used for downstream processing. Lysates were centrifuged at 5,000 × g for 10 minutes, and the supernatants were collected. Proteins were reduced with 10 µL of 200 mM dithiothreitol (Sigma Aldrich, D9779) for 1 h at 57°C with shaking at 700 rpm. Samples were subsequently alkylated with 15 µL of 200 mM iodoacetamide (Sigma Aldrich, I6125) for 1 h at room temperature in the dark. Proteins were digested overnight at 37°C using trypsin/LysC (Promega, V5073) at a ratio of 1 µg enzyme per 30 µg protein. Digestion was terminated by addition of 15 µL of 20% formic acid. Samples were centrifuged and passed through a 10-kDa molecular-weight cutoff filter (Thermo Fisher Scientific, 88513) to remove undigested proteins. Peptides were desalted using C18 spin columns (Thermo Fisher Scientific, 89870) according to the manufacturer’s instructions in low protein binding collection tubes (Thermo Fisher, 90410), dried in a vacuum concentrator, and resuspended in 7 µL of 0.3% formic acid (Thermo Fisher, A117-50). Approximately 1 µg of peptides was injected for LC–MS/MS analysis.

#### LC–MS/MS data acquisition

Proteomic analysis was performed using a Q Exactive HF mass spectrometer (Thermo Fisher Scientific) coupled to an UltiMate 3000 RSLCnano system. Full MS spectra were acquired over an m/z range of 375–1600 at a resolution of 60,000, with a maximum injection time of 60 ms and an AGC target of 1 × 10⁶. The 25 most intense precursor ions were selected for fragmentation.

MS/MS spectra were acquired at a resolution of 15,000 with an AGC target of 1 × 10⁵, a maximum injection time of 100 ms and an isolation window of 1.3 m/z. The first mass was set to 100 m/z. Peptide separation was performed using a nonlinear 180-min gradient at a flow rate of 0.300 µL/min, using 0.1% formic acid in water as solvent A and 80% acetonitrile containing 0.1% formic acid as solvent B.

#### Proteomic data processing

Raw LC–MS/MS data were processed using Proteome Discoverer v2.5. Protein identification was performed against the UniProtKB *Capra hircus* database (UP000291000) together with the sheep database (UP000002356). Carbamidomethylation of cysteine residues was used as a fixed modification and methionine oxidation as a variable modification. Peptide lengths between 6 and 144 amino acids and up to two missed cleavages were permitted. Precursor and fragment mass tolerances were set to 10 ppm and 0.02 Da, respectively. False-discovery-rate filtering was applied as specified in the original analysis. Minora Feature Detector, Precursor Ions Quantifier and Feature Mapper nodes were used for label-free quantification. Protein abundance values were log₂-transformed before statistical analysis. PCA, PLS-DA, hierarchical clustering, volcano-plot analysis and functional enrichment were performed using GraphPad Prism, MetaboAnalyst 6.0 and FunRich 3.1 as appropriate.

#### Proteome and Metabolome raw data

Proteome data submitted to Massive: Massive ID: MSV000103828 Password: V4x850b547vwm0Mr

Metabolome data deposited in Mendeley https://data.mendeley.com/preview/fsmdv62pj6?a=d8da00e1-d10c-42ce-a88c-2ef24f2d3d92

#### Fluorescence-based live-cell imaging

Live-cell fluorescence assays were performed on denuded oocytes to assess intracellular ROS, calcium homeostasis, mitochondrial state, and ER-associated fluorescence.

#### Measurement of intracellular ROS

Reactive oxygen species were measured using 2′,7′-Dichlorodihydrofluorescein diacetate (H₂DCFDA) (Invitrogen, D399). Excellent- and poor-grade oocytes were incubated with 10 µM H₂DCFDA in TCM-199 for 5 min at 37°C. Oocytes were subsequently washed with fresh TCM-199 to remove excess probe and imaged using a Zeiss Axio Observer Z1/7 fluorescence microscope with AF488 detection. Fluorescence intensity was quantified using ZEN software.

### Measurement of cytosolic Ca^2+^

Cytosolic Ca²⁺ was measured using Fluo-3 AM (Invitrogen, F1242). A 500 µM stock solution was prepared in DMSO and diluted to a final concentration of 5 µM in maturation medium. Oocytes were incubated in 100 µL of the working solution for 40 min at 37°C in a CO₂ incubator. Following staining, oocytes were washed three times with fresh TCM-199 and imaged using AF488. Fluorescence intensity was quantified for individual oocytes.

#### Measurement of mitochondrial Ca^2+^

Mitochondrial Ca²⁺ was assessed using Rhod-2 AM (Invitrogen, R1245MP). A 500 µM stock solution was prepared in DMSO and diluted to 5 µM in maturation medium. Oocytes were incubated in 100 µL of the working solution for 40 min at 37°C. Following three washes with fresh medium, fluorescence was imaged using AF555 and quantified.

#### Assessment of mitochondrial membrane potential

Mitochondrial membrane potential was assessed using JC-1 (Invitrogen, M34152). A 100 µM stock solution was prepared in DMSO, and oocytes were incubated with 2 µM JC-1 for 30 min at 37°C. After three washes with TCM-199, fluorescence was immediately recorded using 488-nm excitation for JC-1 monomers and 555-nm excitation for JC-1 aggregates. The red-to-green fluorescence ratio was used as the measure of mitochondrial membrane potential.

#### Mitochondrial staining

Mitochondrial content and distribution were assessed using MitoTracker Green (Invitrogen, M7514). A 5 µM stock solution was prepared in DMSO and diluted to 200 nM in maturation medium immediately before use. Oocytes were incubated with the working solution for 20 min at 37°C and washed three times with TCM-199. Fluorescence was recorded using AF488.

#### ER staining

ER-associated fluorescence was assessed using ER-Tracker Green (Invitrogen, E34251). A 1 mM stock solution was prepared in DMSO and diluted to 5 µM in 100 µL of maturation medium. Oocytes were incubated for 20 min at 37°C and subsequently washed three times with TCM-199. Fluorescence images were acquired using AF488, and fluorescence intensity was quantified.

For all fluorescence-based assays, image acquisition and quantitative analysis were performed using the same imaging settings within each experiment.

#### Immunofluorescence analysis

For immunofluorescence analysis, zona pellucida was removed using 0.2% pronase (Sigma Aldrich, P8811). Oocytes were fixed with 4% paraformaldehyde (PFA) (Sigma Aldrich, 158127) for 30 min at room temperature and washed with 0.05% TBST. Samples were permeabilized with 0.25% Triton X-100 (SRL, 64518) for 15 min at room temperature.

For detection of 5-methylcytosine (5mC), oocytes were denatured with 4 N Hydrochloric Acid (HCl, Sigma, 30721) for 10 min at room temperature and immediately neutralized with 100 mM Tris-HCl (SRL, 16622), pH 8.5, for 10 min. Oocytes were refixed with 4% PFA for 20 min and washed three times with 0.05% TBST. Following an additional permeabilization step, oocytes were blocked in antibody dilution buffer (ADB, 3% BSA, 0.05% Triton X 100, 1XTBS, 10% normal goat serum) for 1 h, with the blocking solution changed every 30 min.

Oocytes were incubated overnight at 4°C with primary antibodies against 5mC (Cell Signaling Technology, CST, D3S2Z), and α-tubulin (Sigma Aldrich, F2168) at a dilution of 1:200 in ADB. After three washes with 0.05% TBST, samples were incubated for 1 h at 37°C with the appropriate fluorophore-conjugated secondary antibody at 1:2000. Oocytes were washed three times with 0.5% TBST and mounted using ProLong Diamond Antifade Mountant containing DAPI (Invitrogen, P36970).

For α-tubulin staining, the acid-denaturation and neutralization steps were omitted. A directly FITC-conjugated anti-α-tubulin antibody (Sigma Aldrich F2893) was used, and therefore no secondary antibody was required.

#### Imaging and quantification Image Acquisition

Depending on the experiment, either Nikon or an Axio Observer Z1/7 fluorescent microscope (Carl Zeiss, Germany) was used to get fluorescence images. The Zeiss Axio Observer Z1/7 microscope was used to obtain images of nuclear signal quantification using a 20× and 10× objective as required, whereas measurements of whole-oocyte fluorescence intensity were obtained using a 10× objective. A 60× objective was used to capture images of fixed oocytes after immunocytochemistry. Apochromat 1.4 NA objectives and EXFO X-Cite metal halide light source.

Images were captured with a Hamamatsu ORCA-ER CCD camera. Every experiment used the same illumination, laser/exposure, detector gain, and acquisition parameters for every image.

### Fluorescence Intensity Analysis

ZEN Blue software (Carl Zeiss, Germany) was used to measure fluorescence intensity. When necessary, Z-stack images were obtained and processed as individual optical sections or maximum-intensity projections. Using the Graphics Tools (Circle or Spline Contour), the whole oocyte was manually delineated as the area of interest (ROI), and each oocyte’s mean fluorescence intensity was considered.

#### RNA isolation and RT-qPCR

Cumulus cells were removed from oocytes before RNA extraction. Oocytes were collected into RNase-free tubes and stored at −80°C until processing. Total RNA was extracted using RNAiso Plus (Takara, 9108). cDNA was synthesized by reverse transcription using a cDNA synthesis kit (Takara, 6110) according manufacture’s. QPRT, ERN1, ATF6, sXBP1, and uXBP1 transcripts target transcript abundance was measured by quantitative real-time PCR using TB Green (Takara, RR820A). pEHIShqprt plasmid was linearized with Sma1 and amplified using Q5 High fidelity DNA Polymerase (New England Biolabs, M0491S).

#### QPRT mRNA stability assay

QPRT transcript stability was assessed using a transcriptional chase with actinomycin D (SRL, 28269). Excellent- and poor-grade oocytes were collected separately, denuded and immediately frozen to establish the 0-h reference samples. A second set of oocytes from each grade was incubated with 10 µg/mL actinomycin D to inhibit transcription.

After 3 h, oocytes were collected, completely denuded, and transferred to RNase-free tubes for storage at −80°C. Total RNA was isolated using RNAiso Plus and QPRT transcript abundance was measured by RT-qPCR. QPRT RNA abundance after 3 h of transcriptional inhibition was compared with the corresponding 0-h level for each oocyte grade. A total of 300 oocytes were used per experimental group.

#### Tunicamycin-induced ER stress and analysis of QPRT expression

To examine the effect of ER stress on QPRT expression, excellent-grade oocytes were cultured in IVM medium containing 5 µg/mL tunicamycin, while untreated excellent-grade oocytes served as controls. Oocytes were collected after 14 h of treatment, denuded and processed for

RNA extraction using TRIzol. QPRT and IRE1α transcript abundance was determined by RT-qPCR. A total of 300 oocytes were used per group.

#### TUDCA treatment and analysis of QPRT expression

To assess whether attenuation of ER stress influenced QPRT expression, poor-grade oocytes were cultured with 200 µM TUDCA (Combi-Blocks, QB-9028) for 14 h. Untreated poor-grade oocytes maintained under identical conditions served as controls. Following treatment, oocytes were denuded and processed for RNA extraction using TRIzol. QPRT and IRE1α transcript abundance was determined by RT-qPCR. A total of 300 oocytes were used per group.

#### In vitro transcription of QPRT RNA

A plasmid containing the human QPRT (pEHIShqprt) gene was obtained from Addgene (992166) and propagated. The plasmid was linearized using SmaI (New England Biolabs, R0141S) in 50ul reaction volume according to the manufacturer and used as a template for PCR amplification. Forward and reverse primers were designed to generate a transcription template containing the T7 promoter and QPRT sequence, with the reverse primer incorporating a poly(A) sequence at the 3′ end.

The amplified DNA template was used for in vitro transcription with the mMESSAGE mMACHINE T7 RNA transcription kit (Thermo Fisher Scientific, AM1344) according to the manufacturer’s instructions. Following transcription, residual DNA template was removed by DNase treatment, and the RNA was purified according to the manufacturer’s protocol.

The pEHIShqprt plasmid was linearized with Sma1 and amplified using Q5 High fidelity DNA Polymerase (New England Biolabs, M0491S).

#### IRE1α-mediated QPRT RNA cleavage assay

The susceptibility of QPRT RNA to IRE1α-mediated cleavage was examined using purified in vitro-transcribed QPRT RNA. Three experimental conditions were analyzed: QPRT RNA incubated with recombinant human IRE1 protein (Abcam, ab271544), QPRT RNA incubated with recombinant human IRE1 protein and STF-083010 (MedChemExpress, HY-15845), and QPRT RNA alone. Equal amounts of 100 ng QPRT RNA were used in all reactions, and all procedures were performed under RNase-free conditions. The purified in vitro transcribed QPRT RNA was subjected to an in vitro cleavage assay with recombinant human IRE1 protein.

Reactions were assembled on ice in RNase-free, low-binding 1.5-mL microcentrifuge tubes in a final volume of 20 μL. Prepared the master mix reaction containing 1 x RNA cleavage buffer and 100 ng QPRT RNA. The 2 x RNA cleavage buffer composition includes 40 mM HEPES (pH 7.4), 200 mM NaCl, 2 mM MgCl₂, 10% glycerol, and 2 mM DTT. The recombinant IRE1 protein was then added to the respective reactions at a final concentration of 100 ng per reaction. Three experimental conditions were prepared: QPRT RNA incubated with recombinant human IRE1 protein, QPRT RNA incubated with recombinant human IRE1 protein in the presence of 30 μM STF-083010, and QPRT RNA incubated with nuclease-free water. The reaction mixtures were incubated for 30 min at 30°C, and the reaction was terminated by the addition of 1 μg proteinase K (Sigma Aldrich, P8044-1G) and incubated at 37°C for 10 minutes, followed by the addition of an equal volume of stop solution, gently mixed, and incubated at 80°C for 5 min and immediately cooled on ice to prevent refolding or reformation of base pairing in the denatured RNA. The stop solution consisted of 10 M urea (SRL, 62762), 0.1% SDS, and 1 mM EDTA (SRL, 40088). Following incubation under the respective reaction conditions, RNA integrity and transcript size distribution were assessed using an Agilent 2100 Bioanalyzer. Electropherograms from IRE1α-treated, inhibitor-treated, and RNA-only samples were compared to determine changes in QPRT RNA integrity.

#### QPRT mRNA microinjection

In vitro-transcribed QPRT RNA was microinjected into the cytoplasm of poor-grade oocytes to determine whether restoration of QPRT expression affected meiotic maturation. Approximately 25 pM QPRT mRNA was injected into each oocyte. Control poor-grade oocytes received an equivalent volume of nuclease-free water using the same microinjection procedure. Following microinjection, oocytes were cultured in IVM medium for 28–32 h and subsequently assessed for meiotic maturation.

#### In silico analysis of QPRT RNA

The full-length goat QPRT transcript, including the 5′ and 3′ untranslated regions, was obtained from the NCBI Nucleotide database using accession XM_018040916.1. QPRT sequences from *Bos taurus*, *Homo sapiens*, *Sus scrofa,* and *Mus musculus* were also retrieved for comparative analysis. The secondary structure of the full-length goat QPRT transcript was predicted using RNAfold from the ViennaRNA package. Minimum-free-energy structures were visualized using Forna to identify predicted stem-loop and potentially single-stranded regions.

The QPRT sequence was screened for previously described IRE1-responsive sequence motifs, including CUG motifs. Candidate motifs located within predicted structured or accessible regions were evaluated as potential IRE1-responsive sites. QPRT sequences from goat, cattle, human, and mouse were aligned using Clustal Omega to assess conservation of candidate regions. RNA accessibility was further evaluated using RNAplfold from the ViennaRNA package. Candidate regions were assessed according to sequence motif, predicted structural accessibility, and evolutionary conservation.

#### Assessment of DNA methylation and hydroxymethylation

Global 5mC levels were assessed by immunofluorescence as described above. Oocytes were fixed, permeabilized, and subjected to acid denaturation before staining with antibodies against 5mC Fluorescence intensity was quantified using identical imaging parameters within each experiment.

#### In vitro fertilization and embryo development

Following IVM, NMN-treated and control excellent-grade oocytes were subjected to in vitro fertilization and subsequent embryo culture to determine whether metabolic rescue of meiotic maturation was associated with improved developmental competence.

Following in vitro maturation (IVM), the matured oocytes were extensively washed in *oocyte and embryo wash media* (WASH, Vitrogen, BO 01) to eliminate any remaining maturation medium and one washed in *in vitro fertilization* media (IVF, Vitrogen, BO 03). *In vitro fertilization* was carried out using frozen Osmanabadi goat semen. The semen was thawed at 36°C before being processed using a Percoll density-gradient centrifugation procedure in accordance with the Vitrogen methodology. In brief, the Percoll gradient was prepared by carefully layering 400 µL of undiluted Percoll conventional (Vitrogen, BO 07), followed by 400 µL of diluted Percoll conventional (200 µL Percoll + 200 µL dilution percol media, BO 06), with 200 µL of thawed semen at the top, maintaining distinct layers without mixing. The gradient was centrifuged at 600 × g for 6 minutes, and the sperm pellet was resuspended in IVF media and centrifuged at 150 × g for 3 minutes. The sperm pellet is resuspended in IVF media and adjusted to 1 million spermatozoa/mL, followed by 5 µL spermatozoa added to 200 µL drop of IVF media with 30-40 oocytes in a 4-well plate (Thermo Fisher, 144444) and co-incubated at 38.5°C in a 5% CO₂ incubator for 20 hours. Following fertilization, presumptive zygotes were washed with IVF medium and subsequently given a final wash in *in vitro culture medium*. The zygotes were then transferred to IVC medium (Vitrogen, BO 04) and cultured for 5 days under standard embryo-culture conditions at 38.5°C in a 5% CO₂ incubator. After five days of in vitro culture, the development of the embryo was evaluated. Embryos were examined, and their developmental stages were recorded based on the number of blastomeres. Embryos that had progressed beyond the 8-cell stage or had attained the 8-cell stage were deemed to have accomplished the defined developmental endpoint. Before being used, IVF media and IVC media were pre-equilibrated with overlaid mineral oil at 37°C under 5% CO₂ in humidified air.

### Analysis of published human oocytes Proteome data

#### Human GV-oocyte proteomic data and sample grouping

Human germinal vesicle (GV)-oocyte proteomic data were obtained from the file GVYOUNG_GVAMA_SPEARMAN.csv, comprising 1,440 protein records measured across 26 oocyte samples. The dataset included 12 young samples and 14 advanced maternal age (AMA) samples. Samples were classified as Young (age ≤30 years) or AMA (age >30 years), using the recorded sample ages and corresponding sample metadata.

#### Principal component analysis

Principal component analysis (PCA) was performed to examine variation in the overall proteomic profiles between Young and AMA oocytes. Proteins with missing measurements in any of the 26 samples were excluded from this PCA. The resulting protein-by-sample matrix was transposed to obtain a sample-by-protein matrix. PCA was performed using centered measurements without scaling each protein to unit variance. The first two principal components were visualized by sample group, with 95% confidence ellipses used to illustrate the group distributions.

#### Targeted analysis of NAD metabolism and ER-stress-associated proteins

A targeted panel of 22 proteins was defined, comprising three NAD-metabolism-associated proteins (NAPRT, NADK2, and SIRT5) and 19 proteins associated with ER stress, protein folding, and proteostasis. UniProt accession identifiers were mapped to human gene symbols using UniProt annotations. Target proteins represented in the measured dataset were extracted for downstream visualization. For the targeted heatmap, protein measurements were standardized within each protein across the analyzed oocytes using row-wise z-score transformation. Samples were displayed in a fixed order, with Young samples followed by AMA samples. Heatmap colors represented relative standardized protein measurements across samples.

#### Comparison of selected proteins between Young and AMA oocytes

Individual boxplots were generated for HSPA5, PDIA3, and SIL1. Each point represented an individual oocyte. Young and AMA groups were compared using Welch’s two-sample t-test. P values were calculated for the three protein comparisons, and the Benjamini–Hochberg procedure was used to adjust P values for multiple testing. A nominal P value <0.05 was considered statistically significant for unadjusted comparisons; FDR-adjusted results were interpreted separately. Figures were generated in R and exported as tiff files at 600 dpi.

### Statistical analysis

Data were analyzed using GraphPad Prism and MetaboAnalyst as appropriate. Comparisons between two experimental groups were performed using two-tailed statistical tests as appropriate for the distribution and experimental design. For metabolomic and proteomic datasets, multivariate analyses, including PCA and OPLS-DA/PLS-DA, were performed as described above. For oocyte-based experiments, a biological replicate rather than the individual oocyte should be considered the experimental unit, where oocytes were pooled within a biological replicate. Data are presented as mean ± SEM or SD unless otherwise specified. Statistical significance was defined according to the thresholds indicated in the corresponding figure legends.

## Data availability

The data that support the findings of this study are available from the corresponding author upon request.

## Methods contact

Further information and requests for resources and reagents should be directed to the lead contact, H.B.D. Prasada Rao

---

**Supplementary Figure 1.**
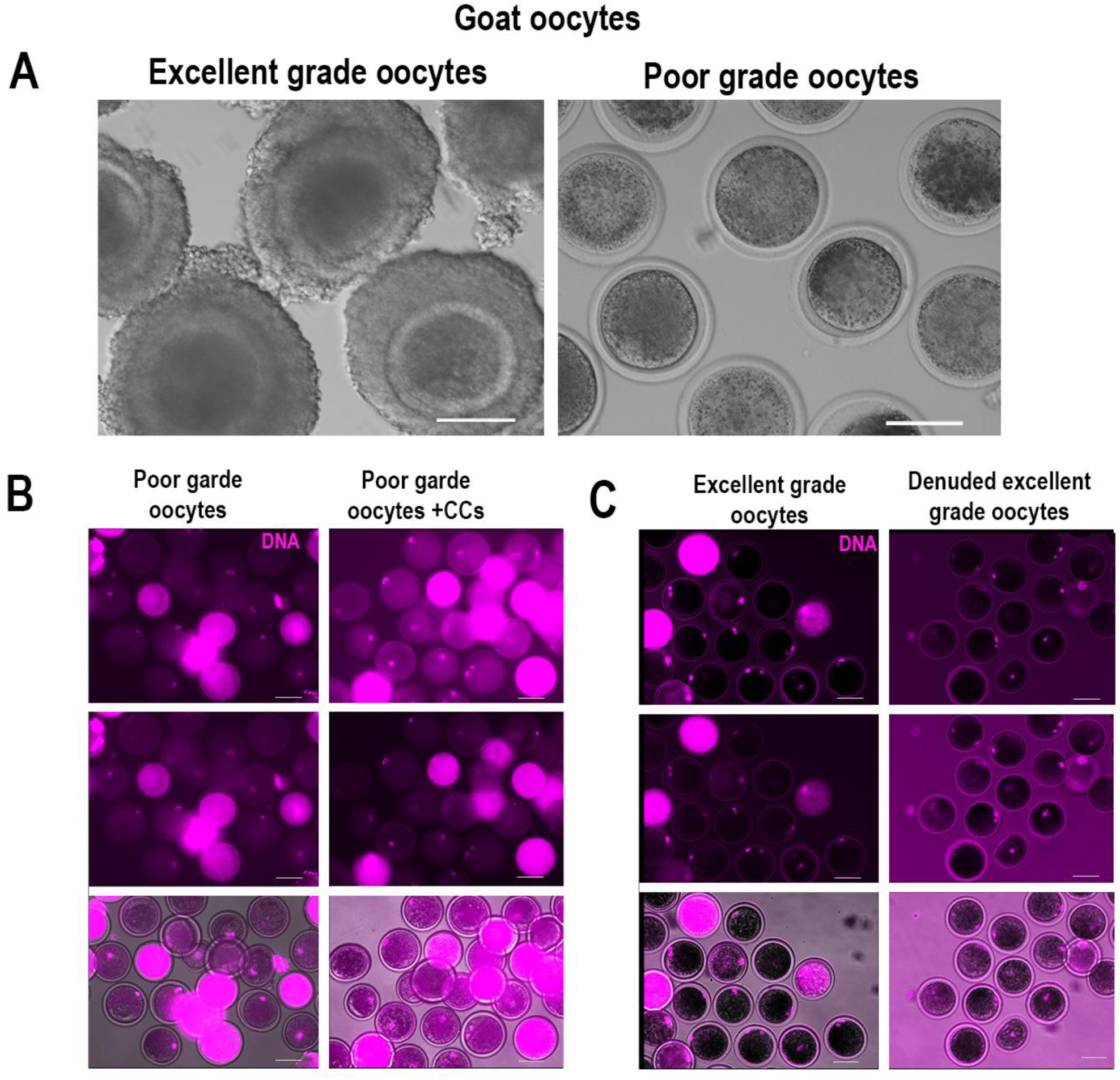
Poor-grade goat oocytes exhibit impaired meiotic competence independent of cumulus cell support. (**A)** Representative bright-field images of excellent-grade and poor-grade goat oocytes based on morphological characteristics. **(B)** Representative images showing the maturation status of poor-grade oocytes and poor-grade oocytes with cumulus cells (CCs) following DNA staining with DAPI (magenta). **(C)** Representative images showing the meiotic maturation stages of excellent-grade oocytes and denuded excellent-grade oocytes following DNA staining with DAPI (magenta). Scale bars: 100 μm.

**Supplementary Figure 2.**
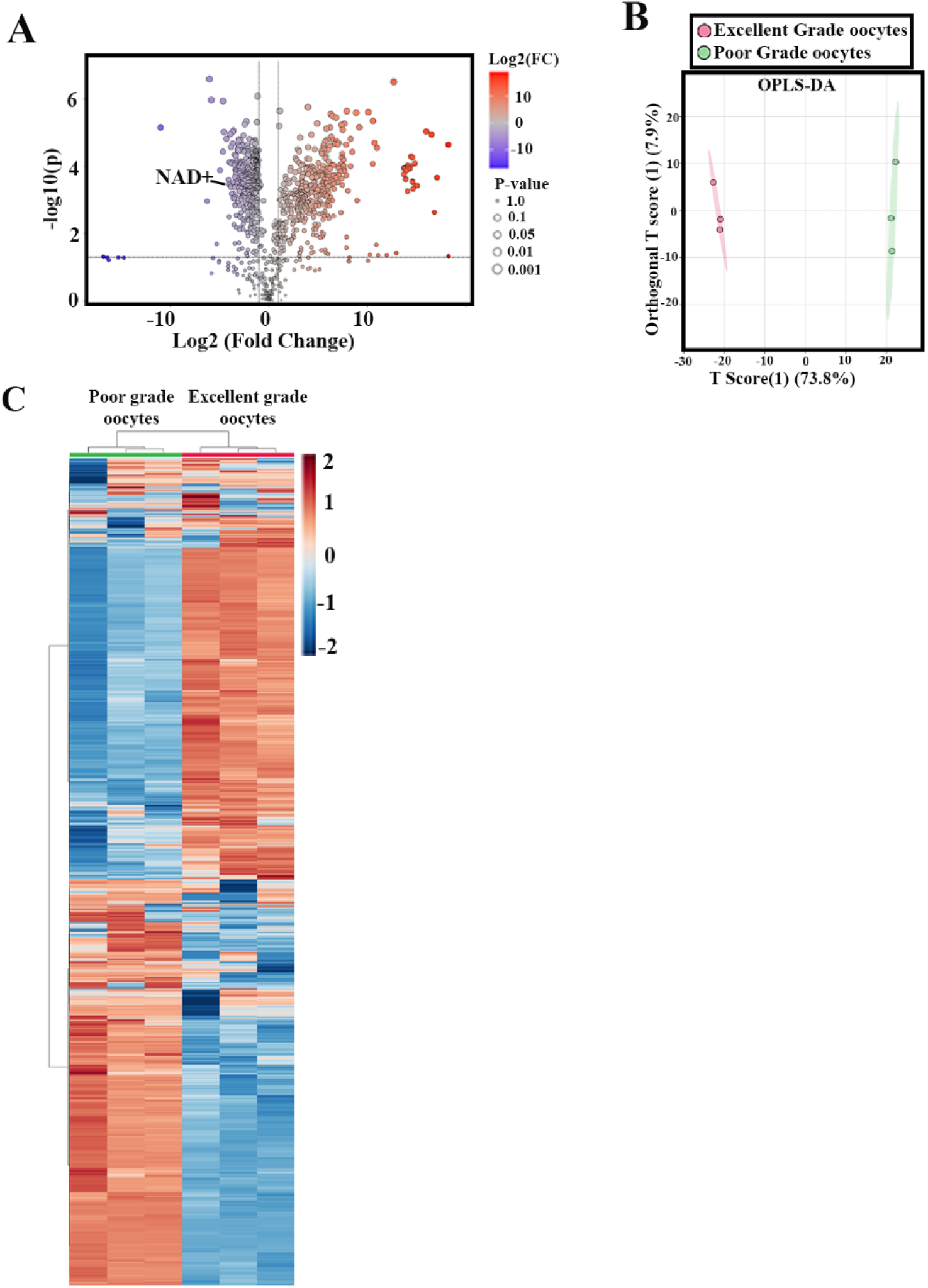
**Metabolomics**(A) Volcano plot showing metabolite fold change between excellent-grade and poor-grade oocytes, highlighting significantly altered metabolites, including NAD+ downregulated in poor-grade oocytes.

**Supplementary Figure 3.**
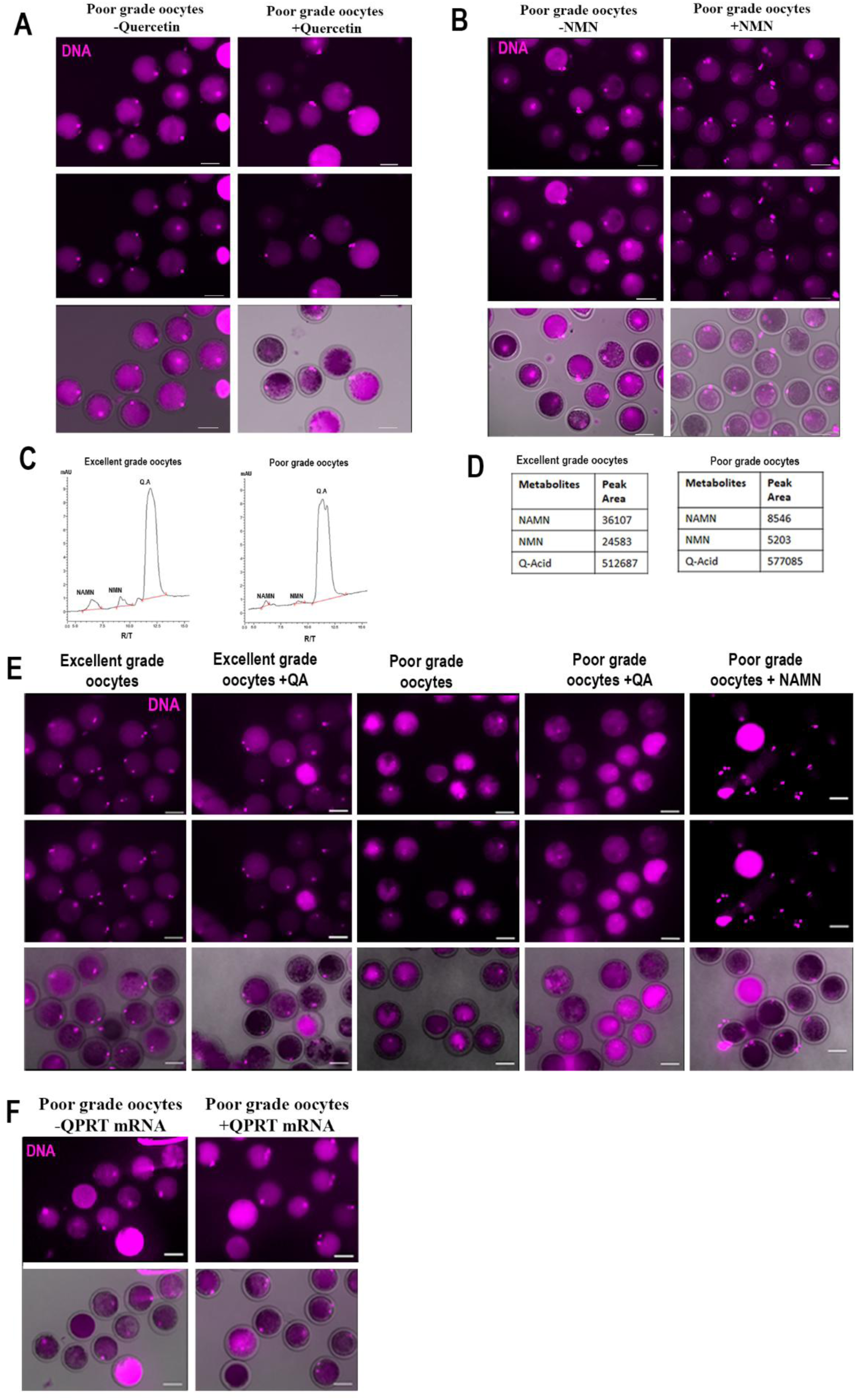
Functional analysis identifies impaired de novo NAD⁺ biosynthesis at the QPRT-dependent step in poor-grade goat oocytes. **(A)** Representative images showing the meiotic maturation of poor-grade oocytes cultured in the absence or presence of quercetin **(B)** Representative images showing the meiotic maturation of poor-grade oocytes cultured in the absence or presence of nicotinamide mononucleotide (NMN). **(C, D)** Representative HPLC chromatograms and peak area showing the detection of nicotinic acid mononucleotide (NAMN), nicotinamide mononucleotide (NMN), and quinolinic acid (Q-Acid) in excellent-grade and poor-grade oocytes. **(E)** Representative images showing the meiotic maturation of excellent grade oocytes with or without quinolinic acid (QA), poor-grade oocytes with or without QA and poor grade oocytes with NAMN following IVM. **(F)** Representative images showing meiotic maturation of poor-grade oocytes following microinjection of QPRT mRNA compared with control oocytes subjected to microinjection without QPRT mRNA. Magenta fluorescence represents DAPI staining of nuclear DNA, and the lower panels show merged fluorescence and bright-field images. Scale bars: 100 μm.

**Supplementary Figure 4.**
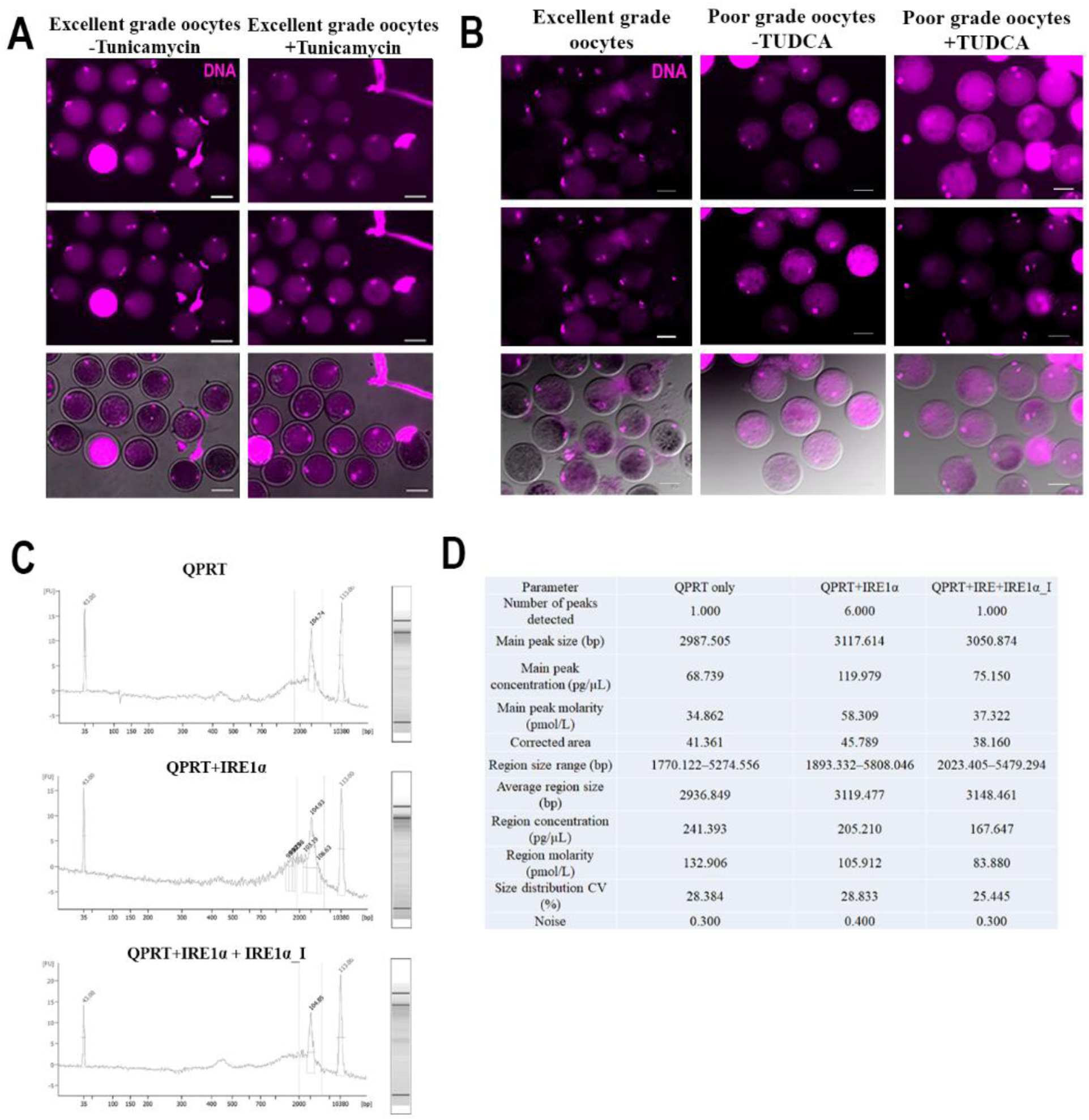
Proteomic profiling identifies ER stress-associated signaling and links IRE1α activation to QPRT dysregulation in poor-grade goat oocytes. **(A)** Representative images showing the meiotic maturation of excellent-grade oocytes treated with or without tunicamycin. **(B)** Representative images showing meiotic maturation of excellent-grade oocytes, poor-grade oocytes with or without TUDCA during in vitro maturation (IVM). **(C)** Representative Agilent 2100 Bioanalyzer electropherograms of QPRT RNA incubated alone, with recombinant IRE1α, and with recombinant IRE1α in the presence of an IRE1α inhibitor (IRE1α_I). The x-axis represents RNA fragment size (bp), and the y-axis represents fluorescence intensity **(D)** Summary table of Bioanalyzer measurements, including the number of detected peaks, main peak size, peak concentration, molarity, corrected area, RNA region size, average region size, region concentration, region molarity, size distribution CV, and noise for the indicated conditions. Magenta fluorescence represents DAPI staining of nuclear DNA. Scale bars: 100 μm

**Supplementary Figure 5.**
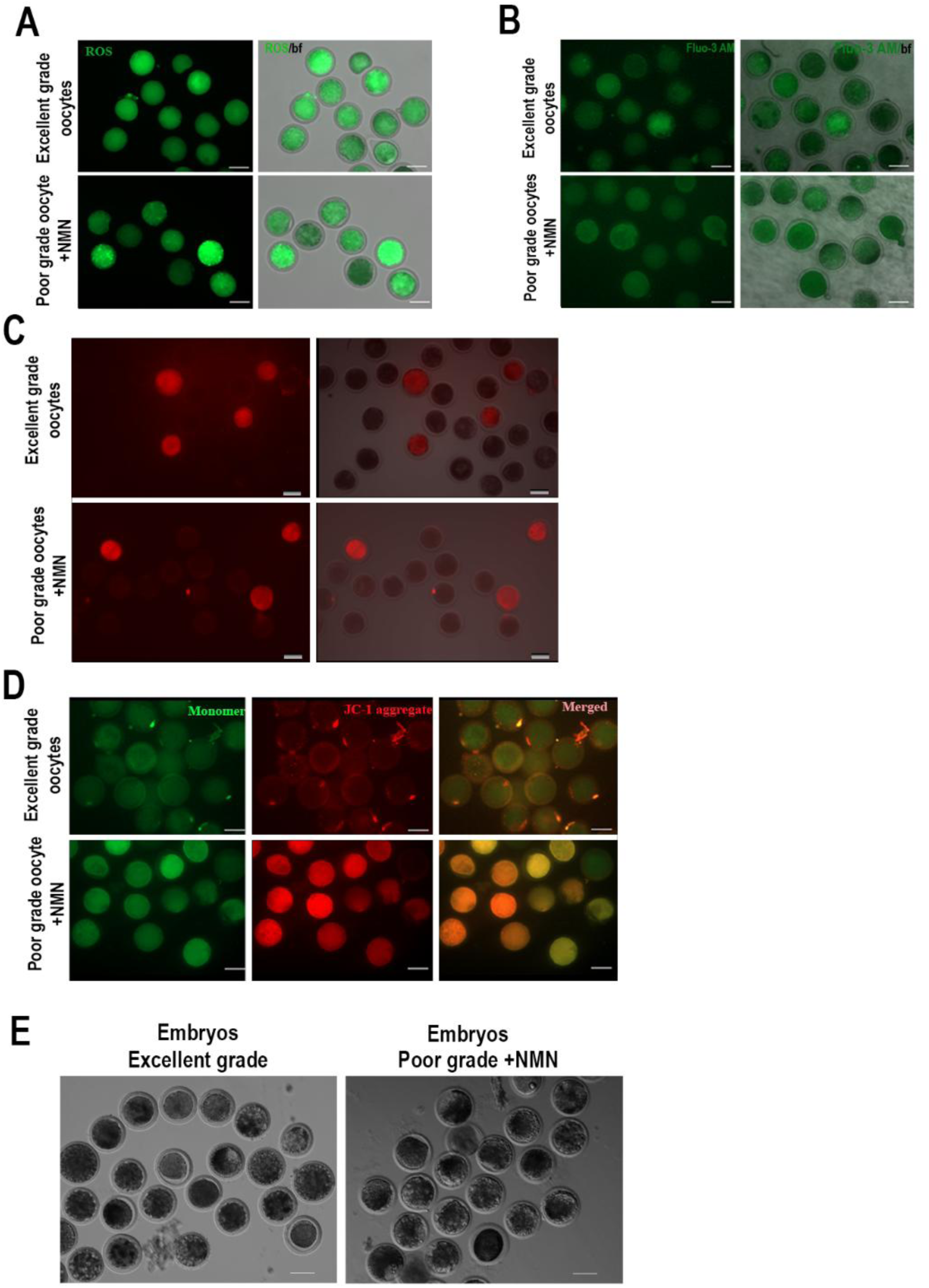
Representative fluorescence images showing NMN-mediated cellular rescue and associated changes in excellent- and poor-grade goat oocytes. **(A)** Representative fluorescence images showing reactive oxygen species (ROS) levels in excellent-grade oocytes and poor-grade oocytes supplemented with nicotinamide mononucleotide (NMN). **(B)** Representative images showing intracellular calcium levels assessed using Fluo-3 AM in excellent-grade oocytes and poor-grade oocytes supplemented with nicotinamide mononucleotide (NMN). **(C)** Representative images showing intracellular mitochondrial Ca²⁺ levels assessed using the Rhod-2 AM in excellent-grade oocytes and poor-grade oocytes supplemented with nicotinamide mononucleotide (NMN). **(D)** Representative fluorescence images showing mitochondrial membrane potential using the JC-1 probe AM in excellent-grade oocytes and poor-grade oocytes supplemented with nicotinamide mononucleotide (NMN), with green fluorescence indicating JC-1 monomers and red fluorescence indicating JC-1 aggregates; merged images show the combined signals. Scale bars: 100 μm.

**Supplementary Figure 6.**
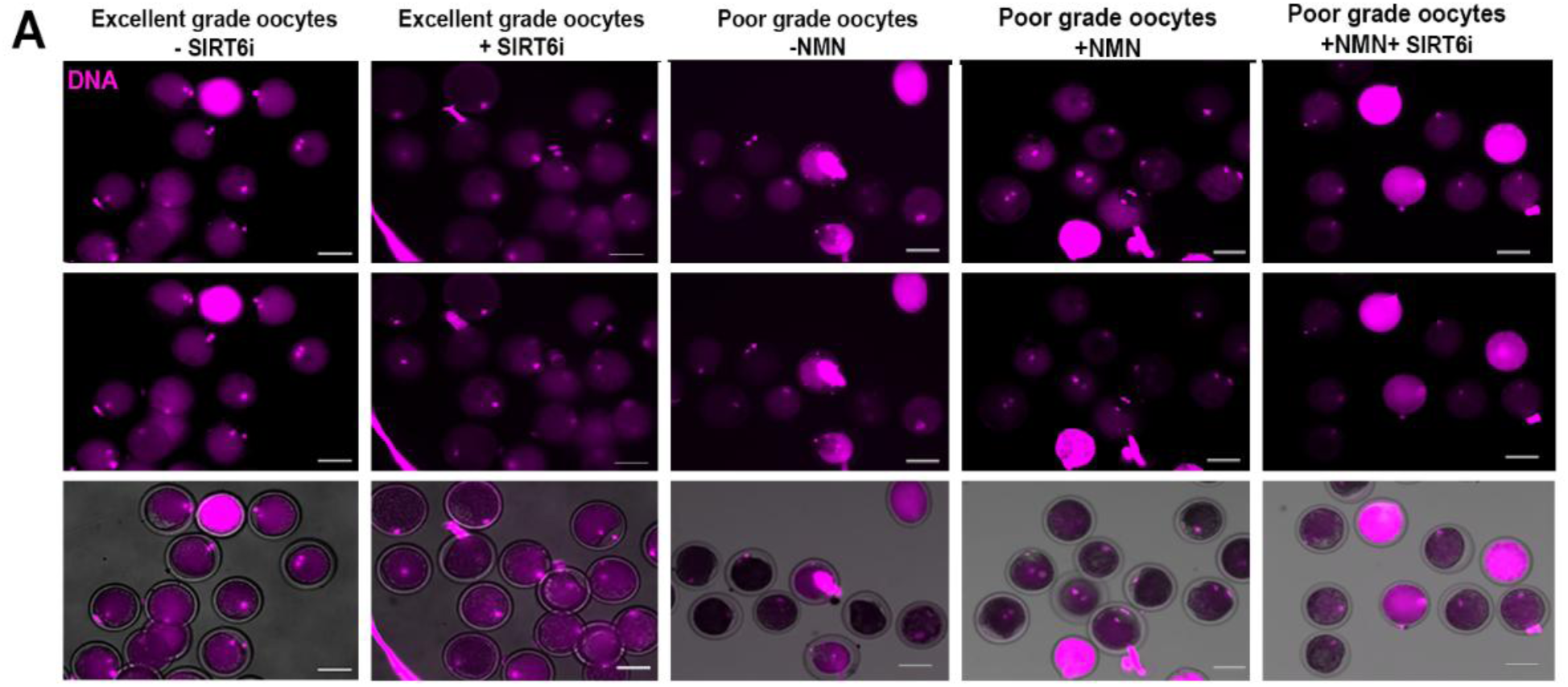
Representative images showing chromatin-associated changes in maturation and SIRT6 dependence in excellent- and poor-grade goat oocytes. **(A**) Representative images showing the meiotic maturation stages of excellent-grade oocytes treated with a SIRT6 inhibitor (SIRT6_I), excellent-grade oocytes supplemented with SIRT6_I, poor-grade oocytes supplemented with nicotinamide mononucleotide (NMN), poor-grade oocytes treated without NMN, and poor-grade oocytes treated with both NMN and SIRT6_i. Magenta fluorescence represents DAPI staining of nuclear DNA. Lower panels show merged fluorescence and bright-field images. Magenta fluorescence represents DAPI staining of nuclear DNA. Scale bars: 100 μm.

**Supplementary Figure 7.**
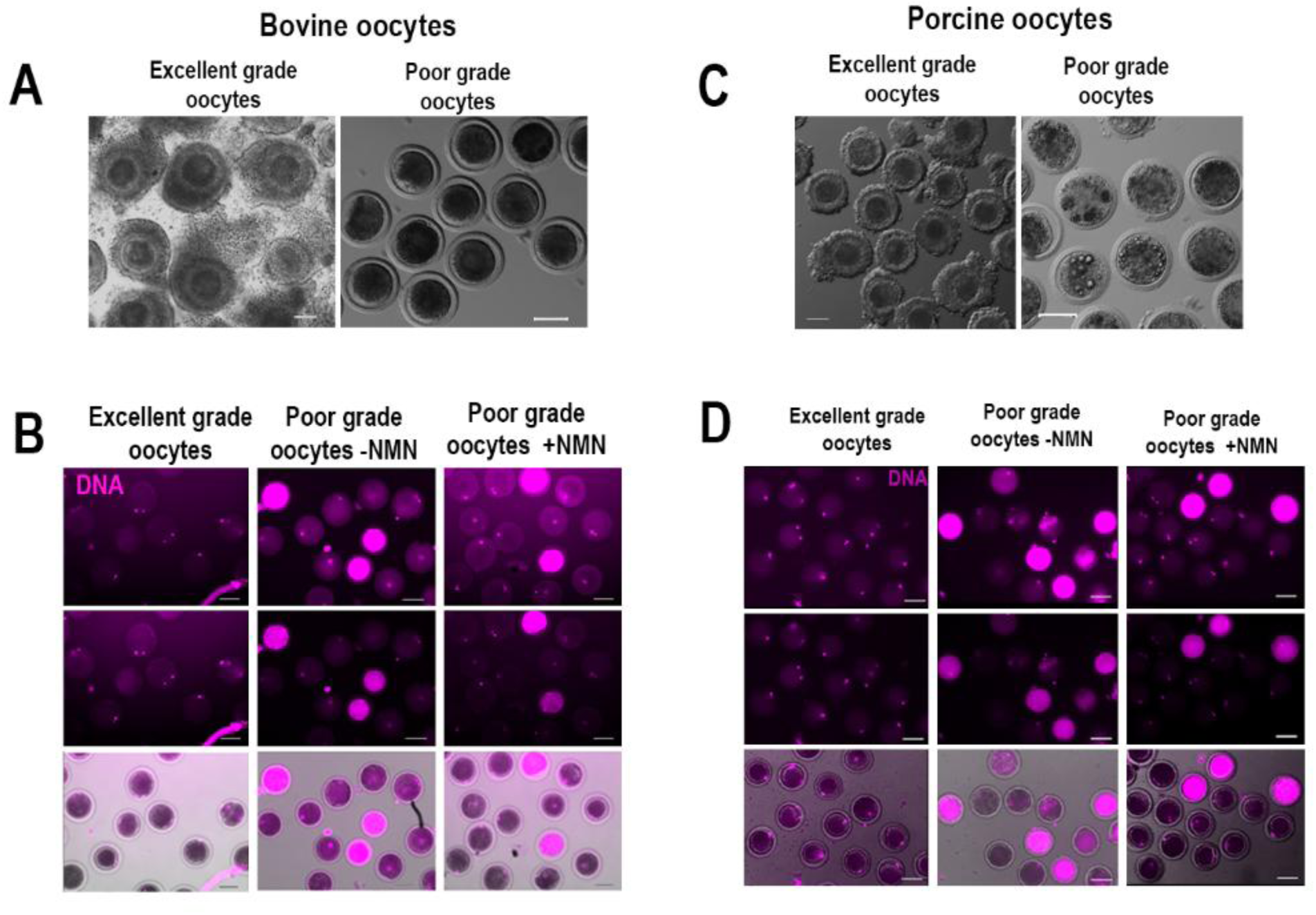
NMN supplementation improves meiotic maturation in poor-grade bovine and porcine oocytes. (**A)** Representative bright-field images of excellent-grade and poor-grade bovine oocytes based on morphological characteristics**. (B)** Representative images showing the meiotic maturation of excellent-grade oocytes, poor-grade oocytes cultured without NMN, and poor-grade oocytes supplemented with nicotinamide mononucleotide (NMN) of bovine oocytes. (**C)** Representative bright-field images of excellent-grade and poor-grade porcine oocytes based on morphological characteristics **(D)** Representative images of excellent-grade oocytes, poor-grade oocytes without NMN supplementation, and poor-grade oocytes supplemented with NMN of porcine oocytes. Magenta fluorescence represents DAPI staining of nuclear DNA, and the lower panels show merged fluorescence and bright-field images. Magenta fluorescence represents DAPI staining of nuclear DNA. Scale bars: 100 μm.

## Model

### ER stress-associated disruption of QPRT-dependent NAD⁺ biosynthesis underlies impaired oocyte quality

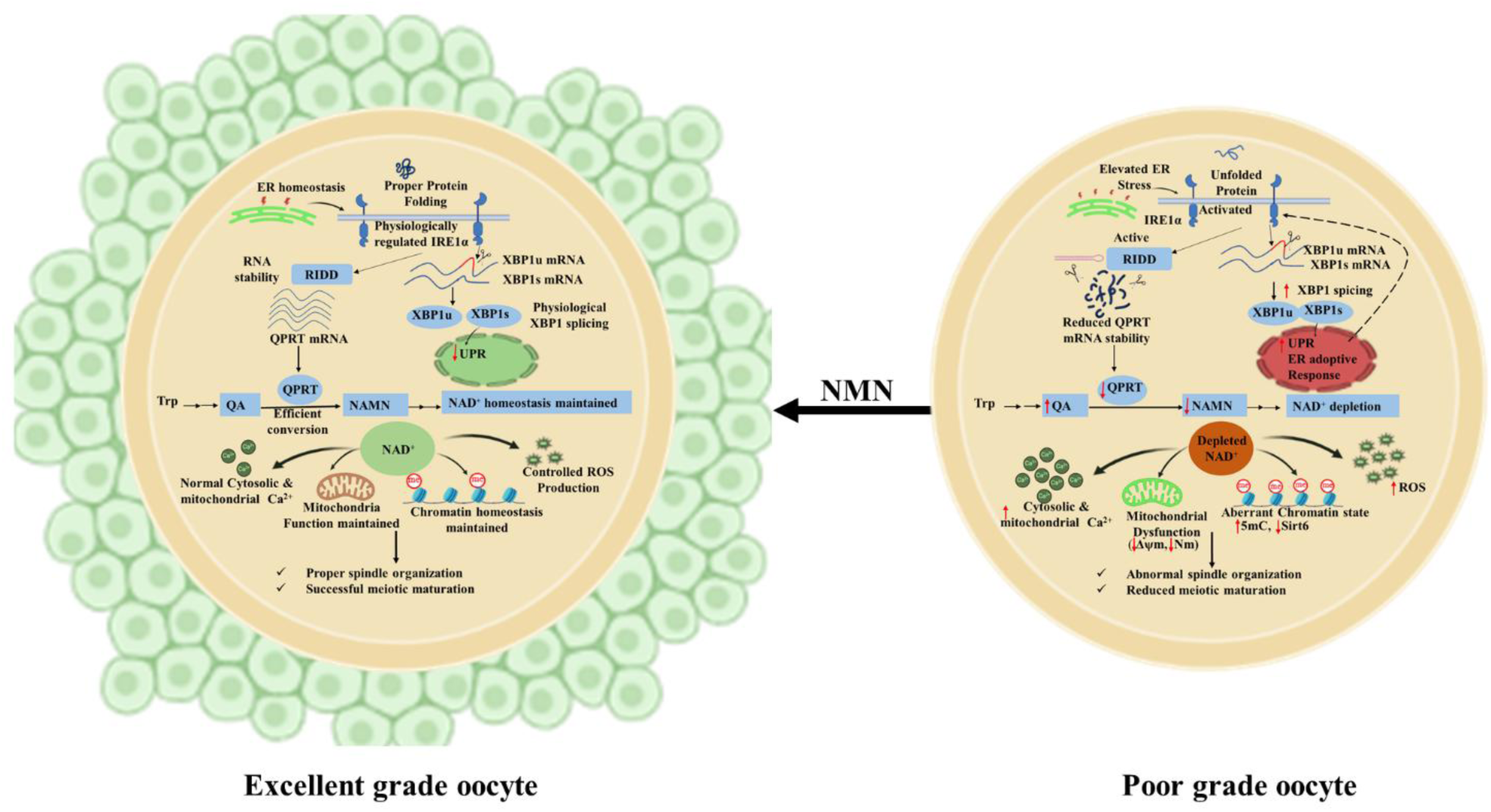

In excellent-grade oocytes, ER homeostasis and IRE1α signalling remain physiologically regulated, supporting the maintenance of QPRT expression and de novo NAD⁺ biosynthesis. Efficient conversion of quinolinic acid (QA) into nicotinic acid mononucleotide (NAMN) helps maintain intracellular NAD⁺ levels. This is accompanied by preserved mitochondrial membrane potential, balanced calcium homeostasis and controlled ROS production. NAD⁺-associated cellular processes, including SIRT6-associated regulation, are also maintained, supporting chromatin organisation, normal spindle formation and successful meiotic maturation.

In contrast, poor-grade oocytes exhibit increased ER stress-associated signalling and IRE1α activation, accompanied by increased XBP1 mRNA splicing and reduced QPRT RNA stability. Our findings suggest that IRE1α may contribute to QPRT RNA processing through a RIDD-like mechanism, resulting in reduced QPRT expression. This impairs the conversion of QA into NAMN, leading to QA accumulation and a marked reduction in NAD⁺ availability. NAD⁺ depletion is associated with mitochondrial dysfunction, reduced mitochondrial membrane potential, increased ROS production and disturbed cytosolic and mitochondrial calcium homeostasis. Alterations in SIRT6-associated regulation and increased global 5mC signal further indicate changes in chromatin-associated processes. Together, these metabolic and cellular disturbances are associated with abnormal spindle organisation, reduced meiotic maturation and impaired support for early embryonic development.

Importantly, supplementation with NMN bypasses the impaired QPRT-dependent step and restores NAD⁺ availability, improving mitochondrial function, calcium homeostasis, oxidative balance and meiotic maturation in poor-grade oocytes.

---

**Table 1:**
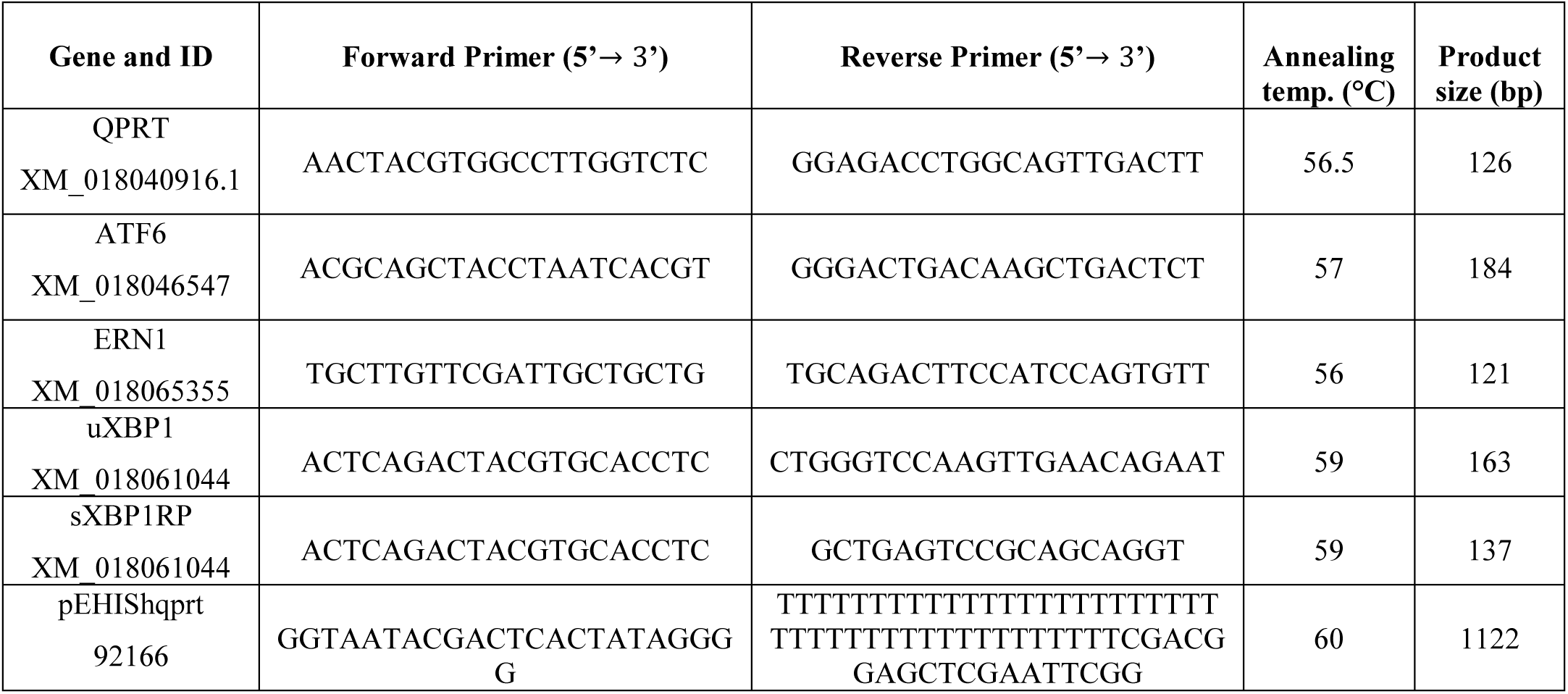
Primer list.

**Table 2:** Inhibitor & enhancer list and concentration.

| **Compound Name** | **Catalogue No.** | **Company** | **Final concentration** |
|---|---|---|---|
| Quercetin | 71923 | SRL | 50 µM |
| NMN | 95130 | SRL | 400 µM |
| L-tryptophan | T0254 | Sigma Aldrich | 750 µM |
| QA | 114941 | Cayman Chemicals | 600 µM |
| NAMN | 32883 | Cayman Chemicals | 400 µM |
| TUDCA | QB-9028 | Combi-Blocks | 200 µM |
| STF-083010 | HY-15845 | MedChemExpress | 30 µM |
| Tunicamycin | 654380 | Calbiochem | 5 µg/mL |
| SIRT6-IN-1 | SML3805 | Sigma, Aldrich | 100 µM |
| Actinomycin D | 28269 | SRL | 10 µg/mL |

**Table 3:** Antibody list.

| **Compound Name** | **Catalogue No.** | **Company** | **Dilution** |
|---|---|---|---|
| 5mC antibody | D3S2Z | CST | 1:200 |
| α-tubulin antibody | F2168 | Sigma Aldrich | 1:200 |

**Table 4:** Dye list.

| Dye Name | Catalogue No. | Company | Concentration | Incubation time (at 37°C) |
|---|---|---|---|---|
| Hoechst 33342 | 14533 | Sigma Aldrich | 2 µg/mL | 10 mins |
| Fmoc-Trp-BODIPY | 900606 | Sigma Aldrich | 20 µM | 40 mins |
| H<sub>2</sub>DCFDA | D399 | Invitrogen | 10 µM | 5 mins |
| Fluo-3 AM | F1242 | Invitrogen | 5 µM | 40 mins |
| Rhod-2 AM | R1245MP | Invitrogen | 5 µM | 40 mins |
| JC-1 | M34152 | Invitrogen | 2 µM | 30 mins |
| MitoTracker Green | M7514 | Invitrogen | 200 nM | 20 mins |
| ER-Tracker Green | E34251 | Invitrogen | 5 µM | 20 mins |

**Table 5:** Reagents and Kits.

| **Reagents/Chemicals** | **Catalogue No.** | **Company** | **Concentration (used)** |
|---|---|---|---|
| Penicillin–Streptomycin | P4333 | Sigma Aldrich | - |
| TCM-199 | M4530 | Sigma Aldrich | - |
| FBS | 10270106 | ThermoFisher | 10% |
| Mineral oil | M8410 | Sigma Aldrich | - |
| LH | L6420 | Sigma Aldrich | 4 IU/mL |
| FSH | F4021 | Sigma Aldrich | 8 IU/mL |
| HEPES | 63732 | SRL | 18 mM |
| sodium pyruvate | P4562 | Sigma Aldrich | 0.81 mM |
| hEGF | GF14 | Sigma Aldrich | 10 ng/mL |
| Gentamicin | G1264 | Sigma Aldrich | 50 µg/mL |
| hMG |  | Sanzyme | 10 IU/mL |
| Hyaluronidase | H3506 | Sigma Aldrich | 0.2 mg/mL |
| DMSO | 66444 | SRL | - |
| Estradiol-17β |  |  | 1 µg/mL |
| Pronase | P8811 | Sigma Aldrich | 0.2% |
| Methanol |  | SRL | - |
| SDC | D6750 | Sigma-Aldrich | 2% |
| AMBIC | 5.33005 | Sigma-Aldrich | 50 mM |
| PFA | 158127 | Sigma-Aldrich | 4% |
| Dithiothreitol | D9779 | Sigma-Aldrich | - |
| Iodoacetamide | 6125 | Sigma-Aldrich | - |
| Trypsin/LysC | V5073 | Promega | 1:30 |
| Triton X-100 | 64518 | SRL | - |
| Hcl | 30721 | Sigma-Aldrich | 4 N |
| Tris-HCl | 16622 | SRL | 100 mM |
| BSA | 83803 | SRL | 3% |
| Normal Goat serum |  |  | 10% |
| ProLong Diamond Antifade Mountant DAPI). | P36970 | Invitrogen | 10 ul |
| Proteinase K | P8044 | Sigma-Aldrich | - |
| recombinant human IRE1 protein | ab271544 | Abcam | - |

**Table 6:** Reagents and Kits.

| **Kits Name** | **Catalogue No.** | **Company** |
|---|---|---|
| NAD kit | MAK037 | Sigma-Aldrich |
| Bradford Protein assay Kit | T9310A | Takara |
| RNAiso Plus | 9108 | Takara |
| cDNA synthesis kit | 6110 | Takara |
| TB Green | RR820A | Takara |
| Q5 High Fidelity Polymerase | M0491S | NEB |
| mMESSAGE mMACHINE T7 RNA transcription kit | AM1344 | Thermo Fisher |
| Sma1 | R0141S | NEB |
| pEHIShqprt | 992166 | Addgene |

